# The impact of genetic modifiers on variation in germline mutation rates within and among human populations

**DOI:** 10.1101/2021.08.25.457718

**Authors:** William R. Milligan, Guy Amster, Guy Sella

## Abstract

Mutation rates and spectra differ among human populations. Here, we examine whether this variation could be explained by evolution at mutation modifiers. To this end, we consider genetic modifier sites at which mutations, “mutator alleles”, increase genome-wide mutation rates and model their evolution under purifying selection due to the additional deleterious mutations that they cause, genetic drift, and demographic processes. We solve the model analytically for a constant population size and characterize how evolution at modifier sites impacts variation in mutation rates within and among populations. We then use simulations to study the effects of modifier sites under a plausible demographic model for Africans and Europeans. When comparing populations that evolve independently, weakly selected modifier sites (2*N*_*e*_*s* ≈ 1), which evolve slowly, contribute the most to variation in mutation rates. In contrast, when populations recently split from a common ancestral population, strongly selected modifier sites (2*N*_*e*_*s* ≫ 1), which evolve rapidly, contribute the most to variation between them. Moreover, a modest number of modifier sites (e.g., 10 per mutation type in the standard classification into 96 types) subject to moderate to strong selection (2*N*_*e*_*s* > 1) could account for the variation in mutation rates observed among human populations. If such modifier sites indeed underlie differences among populations, they should also cause variation in mutation rates within populations and their effects should be detectable in pedigree studies.

## Introduction

Germline mutations are the ultimate source of genetic variation, fueling the evolutionary process. Yet mutation rates themselves also vary and evolve (Elango et al. 2006; Elango et al. 2009; Lynch 2010; Sayres et al. 2011; Scally and Durbin 2012; Ségurel et al. 2014; Harris 2015; Harris and Pritchard 2017; Mathieson and Reich 2017). For example, mutation rates among vertebrates differ by almost an order of magnitude (Kong et al. 2012; Venn et al. 2014; Uchimura et al. 2015; Feng et al. 2017; Lindsay et al. 2019; Harland et al. 2017), and the mutation spectrum differs among primate species (Moorjani et al. 2016; Harris and Pritchard 2017; Goldberg and Harris 2019).

Mutations at loci that modify germline mutation rates, such as in genes involved in DNA replication and repair, can change the rate and spectrum of mutations (Drake 1993; Aarnio et al. 1999; Sasani et al. 2021). Such “mutator alleles” are subject to purifying selection when they increase the burden of deleterious mutations in individuals who carry them (Kimura 1967; Kondrashov 1995; Lynch 2008). Their population frequencies are also affected by genetic drift and demographic processes, such as changes in population size. Therefore, modifiers of germline mutations evolve under an interplay between mutation, purifying selection, genetic drift, and demographic processes, which generates variation in rates and spectrum of germline mutations among individuals, populations, and species.

Differences in the efficacy of selection against mutator alleles could explain variation in mutation rates among evolutionary distant taxa. The strength of selection acting on a mutator allele (quantified by its selection coefficient *s*) should be proportional to its effect on mutation rates (Kimura 1967; Kondrashov 1995; Dawson 1998). Species with effective population sizes, *N*_*e*_, cannot effectively select against mutators with 2*N*_*e*_*s* ≪ 1 (Crow and Kimura 1970) and thus species with smaller *N*_*e*_ are expected to have higher mutation rates (Kimura 1967; Lynch 2008; Lynch 2010; Sung et al. 2012). Variation in *N*_*e*_ may then explain much of the variation in mutation rates among taxa (Lynch 2010; Leffler et al. 2012; Sung et al. 2012) as well as aspects of genome architecture such as variation in genome size (Lynch and Conery 2003; Lynch 2007).

Mutation rates have also been observed to vary substantially over shorter evolutionary timescales, in particular among human populations (Harris 2015; Moorjani et al. 2016; Harris and Pritchard 2017; Mathieson and Reich 2017; Narasimhan et al. 2017; Aikens et al. 2019; Speidel et al. 2019; Goldberg and Harris 2019). When mutations are categorized into 96 different types based on ancestral allele, derived allele, and 5’ and 3’ flanking nucleotides (e.g., GAC→GTC), roughly one-third of these types show significant differences in mutation rates between pairs of continental populations (Harris and Pritchard 2017). Notably, rates of TCC→TTC mutations were estimated to have temporarily increased by 50-100% between 5,000-30,000 years ago in Europeans, whereas these rates have remained approximately constant in Africans (Harris 2015; Harris and Pritchard 2017; Mathieson and Reich 2017; Speidel et al. 2019). Here, we consider how such a large and pulse-like increase in a mutation rate could arise.

Multiple explanations have been proposed for these observations, which are not mutually-exclusive: changes to life history traits that affect mutation rates and spectra, changes in environment that led to differences in mutagen exposures, and evolution at mutation rate modifiers (Harris 2015; Harris and Pritchard 2017; Mathieson and Reich 2017; Carlson et al. 2020; Chintalapati and Moorjani 2020; Macià et al. 2021). Differences in life history traits (notably in generation times) among populations could have generated differences in multiple mutation rates, including the rate of TCC→TTC mutations (Macià et al. 2021). We may be able to learn whether changes in generation times plausibly explain the differences in mutations among populations by testing if these differences align with the effects of parental ages on de novo mutation rates in human pedigree studies (Z. Gao and M. Przeworski, personal communication). Similarly, some environmental mutagens preferentially increase C→T mutation rates, such as alkylating agents that increase G→A mutation rates (Mathieson and Reich 2017) or UV-damage, which has been associated with increased TCN→TTN and CCN→CTN mutation rates in irradiated cells (Drobetsky and Sage 1993; Marionnet et al. 1995) and melanomas (Alexandrov et al. 2020). Additionally, humans populations faced widespread environmental changes during the relevant time period, which may have changed their exposures to such mutagens (Firestone et al. 2007; Broughton and Weitzel 2018; Pino et al. 2019). However, we know little about how relevant exposures changed over time or how various mutagens affect germline mutation rates. Consequently, it is difficult to evaluate whether changes in environment plausibly explain variation in germline mutation rates.

Alternatively, variation in mutation rates among human lineages could reflect evolution at loci that modify germline mutation rates (Harris and Pritchard 2017). Modifier loci that cause variation among populations should also cause variation within populations, which suggests that we may be able to identify their effects in datasets of de novo mutations in pedigrees, genetic variation data from unrelated individuals (see, e.g., Seoighe and Scally 2017), or by quantitative trait locus (QTL) mapping (see, e.g., Sasani et al. 2021). However, to understand whether this explanation is plausible—that is whether evolution at modifier loci could explain observed population differences—requires a model of mutator allele dynamics.

Our goal is therefore to describe how the interplay of mutation, selection, drift, and demographic processes at loci that modify germline mutation rates shapes variation in rates among and within populations. Specifically, we ask: 1) how the number and effect sizes of modifier loci affect their contribution to variation among and within populations; 2) whether evolution at modifier loci could generate the kind of variation observed among human populations, and 3) how we might identify the effects of mutator alleles in datasets of de novo mutations in pedigrees.

### The Model

We assume that an individual’s genotype consists of two parts: *L* strongly selected, biallelic sites that determine fitness and *M* biallelic modifier sites that determine the mutation rate. Specifically, we assume that fitness combines multiplicatively over the selected sites and that each deleterious allele reduces fitness by a factor of (1 − *hs*), where *hs* > 0. Thus, an individual carrying *i* deleterious alleles has absolute fitness (1 − *hs*)^*i*^. Our main results should hold under more general fitness models (e.g., with a distribution of selection effects) and depend primarily on the compound parameter 2*L* · *E*(*hs*). In our mutation model, we assume a baseline mutation rate of *u*_0_ per site per generation, which is the mutation rate obtained with optimized replication and repair machinery (i.e., one that minimizes mutations). We further assume that at each of the *M* modifier sites, one allele is optimal (i.e., it does not increase mutation rates), while the other, mutator allele increases the mutation rate by *ϕ*. Thus, an individual carrying *m* mutator alleles will have mutation rate

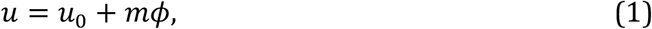

where we assume the effects of mutator alleles are additive across and within sites. Again, our main results should hold under more complex mutation models, e.g., with a distribution of mutator allele effect sizes.

We first model the evolution of mutation rates in a diploid, panmictic population of constant size *N*, with non-overlapping generations. In each generation, individuals are randomly chosen as parents with probabilities proportional to their fitness (i.e., according to Wright-Fisher sampling with fertility selection). These individuals then produce gametes with free recombination among selected and modifier sites, Mendelian segregation, and mutations at both the modifier and selected sites. We assume that the mutational input per generation per selected site is sufficiently low (2*Nu* ≪ 1) and that deleterious alleles are sufficiently rare (2*Nhs* ≫ 1) such that we can apply the infinite sites approximation and ignore reverse mutations. The number of deleterious alleles added per gamete per generation is then Poisson distributed with mean *Lu*. We further assume that the mutation rate at modifier sites per generation, *μ*, is symmetric among alleles and constant (i.e., not affected by mutator alleles). We later show that relaxing this assumption does not affect our qualitative results but may lead to transient surges in mean mutation rates (Fig. S5); we expect this effect will be negligible when individual modifier sites only affect the mutation rate corresponding to a subset of mutation types, as the opportunity for non-linear interactions among modifier sites will be greatly reduced. Finally, we assume that gametes combine randomly to produce the next generation.

We later consider a version of this model under a scenario that mimics a widely-used demographic history inferred for European and African populations by Schiffels and Durbin (2014); hereafter the S&D demographic history (see Fig. S4 for details). Instead of a single, constant-size population, we consider an ancestral population that splits into two independently-evolving populations, where population sizes change over time. For this scenario, we focus on variation among populations in mutation rates affecting one of the 96 potential mutation types, such as TCC→TTC (which have decreased *L* and *M* compared to mutators that affect all mutation types jointly).

### Simulations

We compare our analytic approximations for the mean and variance of mutation rates in a constant-size population with results from simulations that implement this model across a wide range of effect sizes *ϕ* and number of modifier sites *M*. Simulations with large *M* become computationally prohibitive, so we restrict ourselves to simulations with *M* ≤ 10,000. In each simulation, we initialize mutator allele frequencies by sampling from their stationary distribution (see *Results*) and the number of deleterious alleles carried by individuals by sampling from a Poisson distribution with mean 2*Lû*/*hs* (Felsenstein 1974), where *û* is the expected mutation rate (see *Results*). Then, we apply a burn-in period of 10*N* generations such that the population has attained mutation-selection-drift balance at both selected and modifier sites. Each simulation runs for 200N generations. The number of replicates for each combination of parameters is the maximum of 10 and the number needed such that the relative errors in the estimated expectation and variance in mutator allele frequency and expected heterozygosity at modifier sites is less than 25%. We estimate the expectation and variance in mutator allele frequency and the expected heterozygosity at modifier sites based on frequencies sampled every *N* generations and by averaging across all replicates; we calculate heterozygosity assuming Hardy-Weinberg equilibrium (i.e., *π* = 2*q*(1 − *q*)). Standard error in these quantities is calculated as 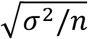, where *σ*^2^ is the variance in the estimated quantity across replicates and *n* is the number of replicates. Increasing the intervals used has little effect on our results. We convert from moments of mutator allele frequencies to moments of mutation rates using simple transformations (Eqs. 13-15).

We also use simulations to investigate whether evolution at modifier sites could account for observed differences in mutation rates between European and African populations. Specifically, we assume the same model as described above with the S&D demographic history of European and African populations (Fig. S4). Here, we separate mutations into 96 different yet equivalent types – corresponding to the 96 types analyzed in previous work (Harris 2015; Harris and Pritchard 2017; Mathieson and Reich 2017; Speidel et al. 2019) – with the rate of each type being affected by its associated *M* modifier sites with effect size *ϕ*. For each mutation type, we simulate mutation rate trajectories – the mean mutation rate in a population recorded each generation – which we later use to assess variation in mean mutation rates among populations.

Generating large samples of mutation rate trajectories for different combinations of *ϕ* and M is computationally prohibitive. Therefore, given *ϕ* and *M*, we generate mutation rate trajectories by sampling *M* allele trajectories from an ensemble of 15,000 allele trajectories at modifier sites with effect size *ϕ*. This ensemble is simulated as follows: 1) We run 15 replicates of a simulation with *M* = 10^3^ modifier sites with effect size *ϕ*, where other parameters take plausible values for humans (see Table 1). 2) The initial mutator allele frequency at each site is sampled from our analytic approximation of the stationary distribution with an effective population size of *N* = 1.4 · 10^4^, the estimate for 7 · 10^4^ generations ago under the S&D demographic history. 3) We run the simulation for a burn-in period of 10^4^ generations with a constant population size of *N* = 1.4 · 10^4^. 4) We simulate evolution at the modifier sites for 6 · 10^4^ generations in the ancestral population followed by 10^4^ generations in each of the two descendant populations separately, where population sizes change according to the S&D demographic history. To generate mutation rate trajectories, we sample *M* of the resulting allelic trajectories, sum their contributions to mean mutation rates in each generation, and add a baseline mutation rate *u*_0_. The *u*_0_ is chosen such that our approximation for the expected mutation rate in a population of constant size *N* = 1.4 · 10^4^ matches a given expected mutation rate *û* (see Table 1). Our simulations exclude the most recent 60 generations, where population sizes are very large. Neglecting this short period should have a negligible effect on our results as there is little opportunity for drift or mutations at modifier sites. Additionally, our results are plausibly insensitive to the minor statistical dependencies among allelic trajectories in our simulations and to our sampling approach more generally (Fig. S10).

**Table 1.** Model parameters and their values used.

| Parameter | Meaning | Value |
| --- | --- | --- |
| $\phi$ | Increase in mutation rate per mutator allele | *Varied* |
| $M$ | Number of modifier sites | *Varied but bounded given $\hat{u}$* |
| $\hat{u}$ | Expected mutation rate at selected sites | $1.25 \cdot 10^{-8}$ or $1.25/3 \cdot 10^{-8}$ |
| $u_0$ | Baseline mutation rate | Fixed given $M, \phi, \hat{u}$ (Eq. 13) |
| $\mu$ | Mutation rate at modifier sites | $1.25 \cdot 10^{-8}$ |
| $Lhs$ | Cumulative fitness effect of mutations at selected sites affected by mutator allele | $1.2 \cdot 10^5$ or $1.2 \cdot 10^5/32$ |
| $N_e$ | Effective population size | $2 \cdot 10^4$ or taken from Schiffels and Durbin (2014) |

### Tests for variation in mutation rates between populations

We use two measures to investigate whether evolution at modifier sites could generate the differences in mutation rates observed between European and African populations. The first mimics the enrichment analysis of Harris and Pritchard (2017), who asked if the fraction of segregating sites corresponding to a given mutation type differed among populations. Specifically, they measured enrichments of a given mutation type in approximately 500 individuals of European ancestry and 500 of African ancestry using the statistic:

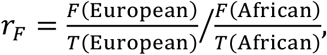

where *F*(*X*) denotes the number of segregating sites corresponding to the focal type in population *X* and *T*(*X*) denotes the total number of segregating sites in population *X*. This statistic is heavily weighted towards recent differences in mutation rates (see Fig. S8).

We mimic their analysis for each combination of *ϕ* and *M*. First, we use msprime (Kelleher et al. 2016) to simulate the ancestral recombination graph (ARG) assuming the S&D demographic history, a sample size of 500 individuals in each population, a genome size of 3 × 10^9^ basis divided into 100 equally sized chromosomes, and a uniform recombination rate of 2 · 10^−8^ per bp per generation. Second, for a given set of 96 mutation rate trajectories (see *Simulations*), we drop mutations corresponding to each mutation type on the ARG and obtain a polymorphism dataset that mimics the one used by Harris and Pritchard (2017) (see SI Section 4). Third, we use this dataset to calculate the enrichment of each mutation type, *r*_*F*_, and record the largest enrichment among the 96. Fourth, we repeat the second and third steps for each of the 10^5^ sets of mutation rate trajectories to obtain an empirical distribution of mutation rate enrichments for a given *ϕ* and *M*. We use the same simulated ARG throughout, but variation in the ARG is negligible (Fig. S7) and its effects on our analysis is largely removed by the ratio for estimating mutation rate differences, F/T.

For our second measure of variation in mutation rates, we consider the average mutation rate across time intervals, which Speidel et al. (2019) estimated for human populations. Specifically, we calculate the average mutation rate in 14 non-overlapping log-sized time intervals along our simulated mutation rate trajectories (seven per population during the 10^4^ generations since they split, with the times demarking the intervals being 60, 124, 258, 537, 1116, 2318, 4814, and 10000 generations). We then test for the peak-like behavior observed in human populations, as follows: 1) We define an interval as *elevated* if its average rate is at least 10% greater than the expected rate or *peaked* if the average is 50% greater than expected. We later relax this definition such that the average rate must be 50% greater than the expected rate for an interval to be elevated (Fig. S11), 2) We classify mutation rate trajectories into three categories: *peak-like* if one interval is peaked and none of the other intervals are elevated; *multi-elevated* if multiple intervals are elevated; or *not-peaked* if none of the intervals are peaked and at most one is elevated. 3) We then classify *a set of mutation rate trajectories* as: *peak-like* if it includes at least one peak-like trajectory but none of the trajectories are multi-elevated; *multi-elevated* if one or more of the trajectories are multi-elevated trajectories; or *not-peaked* if all trajectories are not-peaked. Finally, we estimate the probabilities that a set of mutation rate trajectories would be peak-like, multi-elevated and not-peaked for different values of *M* and *ϕ*.

### Model parameters

The parameters of the model and the values we use for them are detailed in Table 1. In our analyses, we vary the effect size, *ϕ*, and number of modifier sites, *M*, while setting other parameters to plausible values for humans. We also investigate the sensitivity of our results to changes in these parameter values (Fig. S2).

First, consider mutation parameters. Estimates of the average point mutation rate in humans are ∼1.25 · 10^−8^ per site per generation (Kong et al. 2012). Accordingly, we assume that the mutation rate at modifier sites is on average *μ* = 1.25 · 10^−8^ per generation and that the expected mutation rate at selected sites is *E*(*u*) = *û* = 1.25 · 10^−8^ per generation. When we consider a specific mutation type (e.g., *TCC* → *TTC*), we use a third of this *û* estimate. As we detail in *Results*, the expected mutation rate per site given *M* modifier sites with effect size *ϕ* is *û* = *u*_0_ + *M* · 2*E*(*q*)*ϕ*, where *E*(*q*) is the expected mutator allele frequency. We use the estimated value of *u*= to calculate the maximal number of modifier sites with a given effect size *ϕ, M*^∗^(*ϕ*), and to determine the baseline mutation rate *u*_0_ given *M* modifier sites with a given effect size *ϕ*.

Second, consider selection parameters. Selected sites affect mutator allele dynamics through the compound parameter *Lhs*. Estimates of the proportion of the human genome subject to purifying selection range from ∼7-9% (Ponting and Hardison 2011; Ward and Kellis 2012; Kellis et al. 2014; Rands et al. 2014) corresponding to ∼2.1−2.7 × 10^8^ selected sites. We do not currently have good estimates for the average strength of selection at these sites. Assuming that the average selection coefficient is within a factor of two of *E*(*hs*) ≈ 5 · 10^−4^, such that 2*NE*(*hs*) ≈ 20 assuming *N*_*e*_ ≈ 2 · 10^4^ (Schiffels and Durbin 2014). This suggest that genome-wide 2*LE*(*hs*) ≈ 2.4 × 10^5^. We use this value when we consider modifier sites affecting the total mutation rate and 1/32 of this value when we consider the mutation rate at one of 32 potential trinucleotide sequence motifs (e.g., *TCC*). In our analyses, we keep these selection parameters constant (See Table 1) and explore how varying mutator allele effect size *ϕ*, which varies the strength of selection against mutator alleles, impacts their dynamics. Alternative values of parameters yield similar results (Fig. S2).

Lastly, consider the population size. For the model with a constant population size, we use *N*_*e*_ ≈ 2 · 10^4^ (Schiffels and Durbin 2014). For the results corresponding to variation between African and European populations, we use the S&D demographic history (Fig. S4). When we test our analytic predictions for the model with a constant population size against simulations, we use a smaller population size of *N*_*e*_ = 10^3^ for computational efficiency. In this case, we scale other parameters such that their population scaled values, i.e., 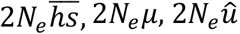, and 2*Lhs*, retain plausible values (i.e., those corresponding to *N*_*e*_ ≈ 2 · 10^4^).

The software used for all simulations, analytic approximations, and figures can be found at https://github.com/sellalab/Mutator.

## Results

### Mutation rates at steady state

We begin by considering the expectation and variance of mutation rates in a population of constant size at steady state. First, we approximate the expected selection coefficient against a mutator allele with a given effect size *ϕ*. Second, we rely on that selection coefficient to derive a diffusion approximation for the stationary distribution of mutator allele frequency. Third, we rely on the stationary distribution to calculate the expectation and variance of the mutation rate in models with many modifier sites and validate our results against simulations. These results provide an understanding of how mutation-selection-drift balance at modifier sites affects mutation rates, which later informs our analysis of the more complicated scenario of variation in mutation rates between African and European populations.

### The expected selection coefficient against a mutator allele

Selection against mutator alleles derives from the excess deleterious germline mutations that individuals with these alleles carry. Kimura (1967) approximated the selection coefficient against mutator alleles based on the rate at which extra deleterious mutations are introduced and the rate at which they are removed by selection, recombination, and Mendelian segregation. Specifically, an individual carrying a single mutator allele produces gametes with an expected *Lϕ* extra *de-novo* deleterious mutations. Assuming free recombination and Mendelian segregation, we expect half of the extra mutations transmitted with the mutator allele in the previous generation to be transmitted with it to the next generation. Under these assumptions, the number of extra mutations that are linked to a mutator alleles that arose *t* generations ago, *X*(*t*), is Poisson distributed with mean

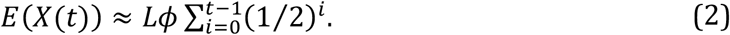

Each mutation stays linked to the mutator allele for two generations on average, so this mean number quickly approaches twice the expected difference in *de novo* mutations, *E*(*X*(∞)) = 2*Lϕ*. Additionally, each mutation carries a multiplicative fitness loss of (1 − *hs*). When mutator alleles are sufficiently old (e.g., *t* > 10), then the expected relative fitness of an individual carrying a mutator allele is

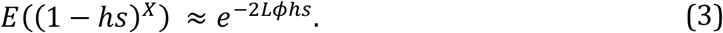

suggesting that the expected selection coefficient against mutator alleles is approximately 1 − *e*^−2*Lϕhs*^.

This approximation becomes inaccurate when selection is very weak or very strong. When selection is weak, mutator alleles can reach high frequencies. High frequency alleles reduce the population’s average fitness and thus experience weaker selection, yet they may be found as homozygotes, which experience stronger selection. The combination of these effects reduces the strength of selection by a factor 1 − *q* (SI Section 2). When selection is very strong, mutator alleles experience weaker selection for a different reason: they do not persist long enough to accumulate 2*Lϕ* extra mutations. For this case, we calculate the expected strength of selection against mutator alleles, *s*^∗^, accounting for their short lifespans (SI Section 1). We combine both corrections and approximate the selection coefficient of mutator alleles by *s*^⋆^(1 − *q*), which is accurate throughout the range of selection effects. Specifically, when selection is strong, *q* ≪ 1 and thus *s*^⋆^(1 − *q*) ≈ *s*^∗^ and when selection is weak, *s*^⋆^ ≈ 2*Lϕhs* (see Fig. S1) and thus *s*^⋆^(1 − *q*) ≈ 2*Lϕhs*(1 − *q*).

### The stationary distribution of mutator alleles

Given the selection coefficient against mutator alleles, we calculate the stationary frequency distribution at the bi-allelic modifier sites. We do so using the diffusion approximation based on the first two moments of change in mutator allele frequency in a single generation (Crow and Kimura 1970; Ewens 2004). We separate the 1^st^ moment into the expected frequency change due to mutation

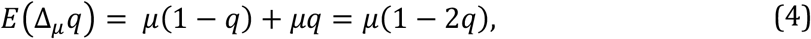

where the two terms correspond to mutations to and from the mutator allele, respectively, and the expected change due to selection

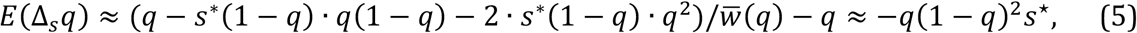

where the two middle terms correspond to selection against heterozygotes and homozygotes for the mutator allele, respectively, and 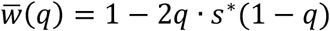 is the relative mean fitness. Taken together, we approximate the expected change in mutator allele frequency by

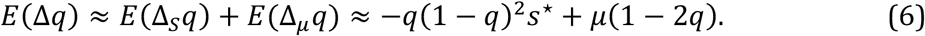

In turn, the variance of change in mutator allele frequency (the 2^nd^ moment) is well approximated by the standard drift term:

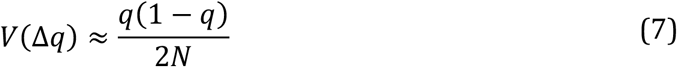

under the standard assumption that *s*^⋆^ ≪ 1. Based on these moments, the stationary distribution takes the form:

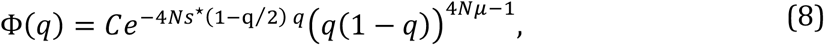

where *C* is a normalizing constant such that 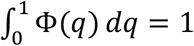 (Fisher 1923; Wright 1945; Kimura 1955; Ewens 2004). Since 4*Nμ* ≪ 1, the integral of the distribution is singular at the boundaries; therefore, we use a standard approach to discretize Φ(*q*) near 0 and 1 in order to calculate summaries of interest (Ewens 2004, SI Section 3). This distribution depends only on the two compound parameters: the scaled mutation rate 4*Nμ* and the scaled selection parameter 4*Ns*^⋆^. In the analyses below, we vary the effect size *ϕ*, which in turn changes 4*Ns*^∗^ and the strength of selection against mutator alleles. We measure the strength of selection in terms of the scaled selection parameter 4*NhsLϕ*.

### Expectation and variance of the mutation rate

We use the stationary distribution to approximate the effect of modifier sites on the expectation and variance of the mutation rate. We first consider the effects of a single modifier site. The expected increase in the mutation rate is given by

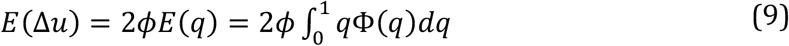

We use the law of total variance to divide the variance in mutation rates into expected variance in mutation rates within a population, *E*(*V*(*u*|*q*)), and the variance of the mean mutation rate of a population, *V*(*E*(*u*|*q*)), which reflects variance between populations, such that

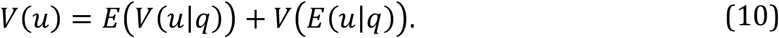

The variance within a population is

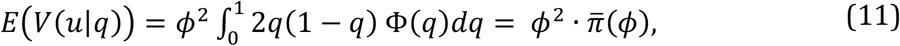

where 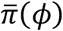 is the expected heterozygosity at a modifier site with effect size *ϕ*. The variance between populations is

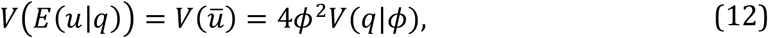

where *ū* is the mean rate in the population with a given mutator allele frequency, which we refer to as the *mean mutation rate*. Most of our analyses focus on variance between populations, *V*(*ū*), but we return to the expected variance within a population in the *Discussion*. Assuming linkage equilibrium (LE) among modifier sites, the expressions for the expectation and variance of mutation rates given *M* modifier sites become:

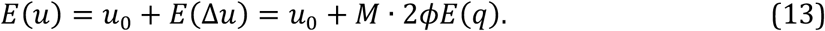

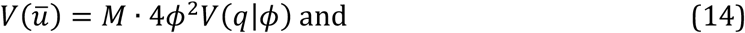

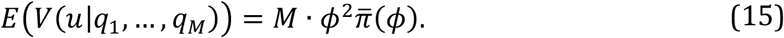

Our approximations perform well throughout the range of selection regimes and a range of numbers of modifier sites when compared to simulation results (Fig. 1).

**Figure 1.**
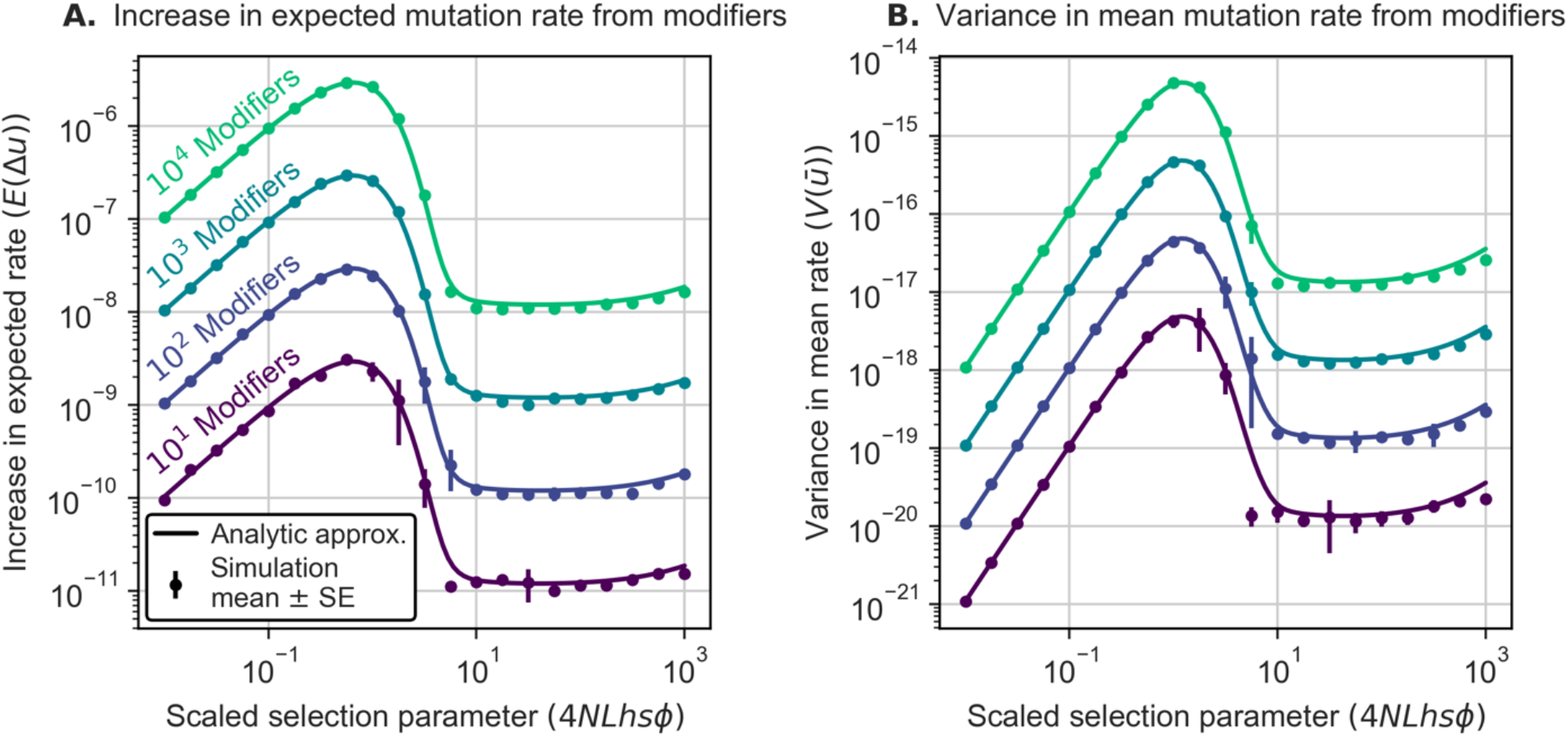
Effect of modifier sites on (A) expected mutation rates and (B) variance of mean mutation rates. We estimate both quantities and their standard error (SE) for each combination of *M* and the scaled selection parameter 4*NLhsϕ* (varying *ϕ*) from simulations (see *Simulations*). In most cases, the SEs are too small to see and therefore not shown. Analytical results are calculated from Eqs. 13 and 14. We assume *N* = 10^3^ with other parameters chosen to match population scaled values in humans (see *Parameters* and Table 1).

The different regimes of modifier dynamics reflect a trade-off between effect size and frequency. A similar trade-off underlies the dependence of genetic load on the selection coefficient (Haldane 1937; Kimura et al. 1963; Kondrashov 1988; Simons et al. 2014). When mutator alleles are effectively neutral (2*Ns*^⋆^ ≈ 4*NLhsϕ* ≪ 1), selection is too weak to affect allele frequencies. Expected mutation rates therefore increase linearly with *ϕ* and variance in mean rates increase linearly with *ϕ*^2^ (Eqs. 13 and 14); both increases arise primarily from substitutions at modifier sites, half of which are fixed for the mutator allele, rather than from segregating mutator alleles (as 4*Nμ* ≪ 1). In the nearly neutral regime (2*Ns*^⋆^ ≈ 4*NLhsϕ*∼1), selection against mutator alleles becomes effective, and fewer sites are fixed for mutator alleles. Effect sizes, *ϕ*, in this regime cause the greatest increase in expected rates and variance in mean rates. When selection is strong (2*Ns*^⋆^ ≈ 4*NLhsϕ* ≫ 1), modifier sites follow the classic mutation-selection balance model and mutator alleles are restricted to low frequencies. Increases in effect size are countered by a proportional decrease in frequency yielding approximately constant expected rates and variance in population mean rates (Eqs. 13 and 14). Lastly, when selection becomes very strong (e.g., *s*^⋆^ > 0.1), *s*^⋆^ is substantially smaller than the selection parameter 2*hsLϕ* (Fig. S1), and increases in effect size are not fully countered by decreases in frequency, resulting in an increase of the expected rates and variance of population mean rates.

Considering the trade-off between effect size and frequency also clarifies how factors we did not include in the model would affect the results. For instance, we assume free recombination among sites. Linkage disequilibrium (LD) between modifier and selected sites would increase the excess number of deleterious mutations linked to mutator alleles relative to the number in our model, which would cause stronger selection against mutator alleles (Kimura et al. 1963; Lynch 2008). As a result, mutator alleles would reach lower frequencies and cause smaller increases in the expectation and variance of mutation rates. The strength of selection would still be proportional to effect size (Kimura 1967), so modifier sites would exhibit similar dynamic regimes to those described. Similarly, we only consider effects of modifier sites on germline mutation rates. If some also increase somatic mutation rates, they likely experience additional selection (e.g., because they increase the incidence of cancer) that increases with their effect size (Lynch 2008). Such modifier sites would have reduced effects on germline mutation rates but plausibly exhibit similar dynamic regimes (Fig. S2). In turn, modifier sites that affect only a subset of possible mutation types affect fewer sites (i.e., smaller *L*) compared to less specific modifier sites. When we keep the scaled selection coefficient, 4*NLhsϕ*, constant, more specific modifier sites have a larger effect size, *ϕ*, resulting in a greater expected mutation rate and larger variance at affected sites. However, because the increase in effect size is balanced by a decrease in the number of affected sites, modifier sites matched for 4*NLhsϕ* result in the same increase in the expectation and variance of the total mutation rate.

### Bounds on the number of modifier sites

Our results suggest that the expectation and variance of mutation rates are proportional to the number of modifier sites (Eqs. 13-15). Investigating the potential variance in mutation rates therefore requires placing bounds on the potential number of modifier sites. One approach is to consider the number and size of genes involved in DNA replication and repair. In humans, there are 742 unique genes under the Gene Ontology terms DNA replication (GO:0006260) and DNA repair (GO: 0006281) with an average transcript length of 3Kb, amounting to ∼1.8 million potential modifier sites (Ashburner et al. 2000; The Gene Ontology Consortium 2019). While mutations at many of these sites are likely to be lethal (e.g., those in genes with pLI ≈1 (Lek et al. 2016) or with *hs* ≈ 1 (Cassa et al. 2017)) or to have no effect on mutation rates (e.g., synonymous), some regulatory sites will affect mutation rates as well. These considerations suggest that there could be as many as hundreds of thousands of modifier sites.

As an alternative approach, we bound the potential number of modifier sites by requiring the expected mutation rate *E*(*u*) to match the total mutation rate estimate of 1.25 × 10^−8^ per bp per generation in humans. When we consider modifier sites that affect only a given mutation type (e.g., *TCC* → *TTC*), we take the expected rate to be a third of this estimate. If we assume the estimated rate *û* is close to the expectation *E*(*u*) and arises entirely from mutation-selection-drift balance at modifier sites (i.e., the baseline rate *u*_0_ = 0), rearranging Eq. 13 suggests that the maximal number of modifier sites with effect size *ϕ* is

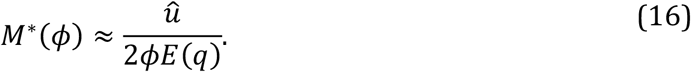

Substituting *M*^∗^(*ϕ*) into Eq. 14, we find that the corresponding variance in mean mutation rates is

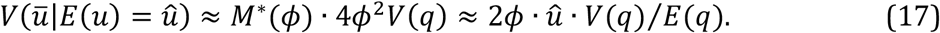

For brevity, we denote this variance by *V*(*ū*|*û*). These bounds on the numbers of modifier sites and corresponding variances are shown in Fig. 2.

**Figure 2.**
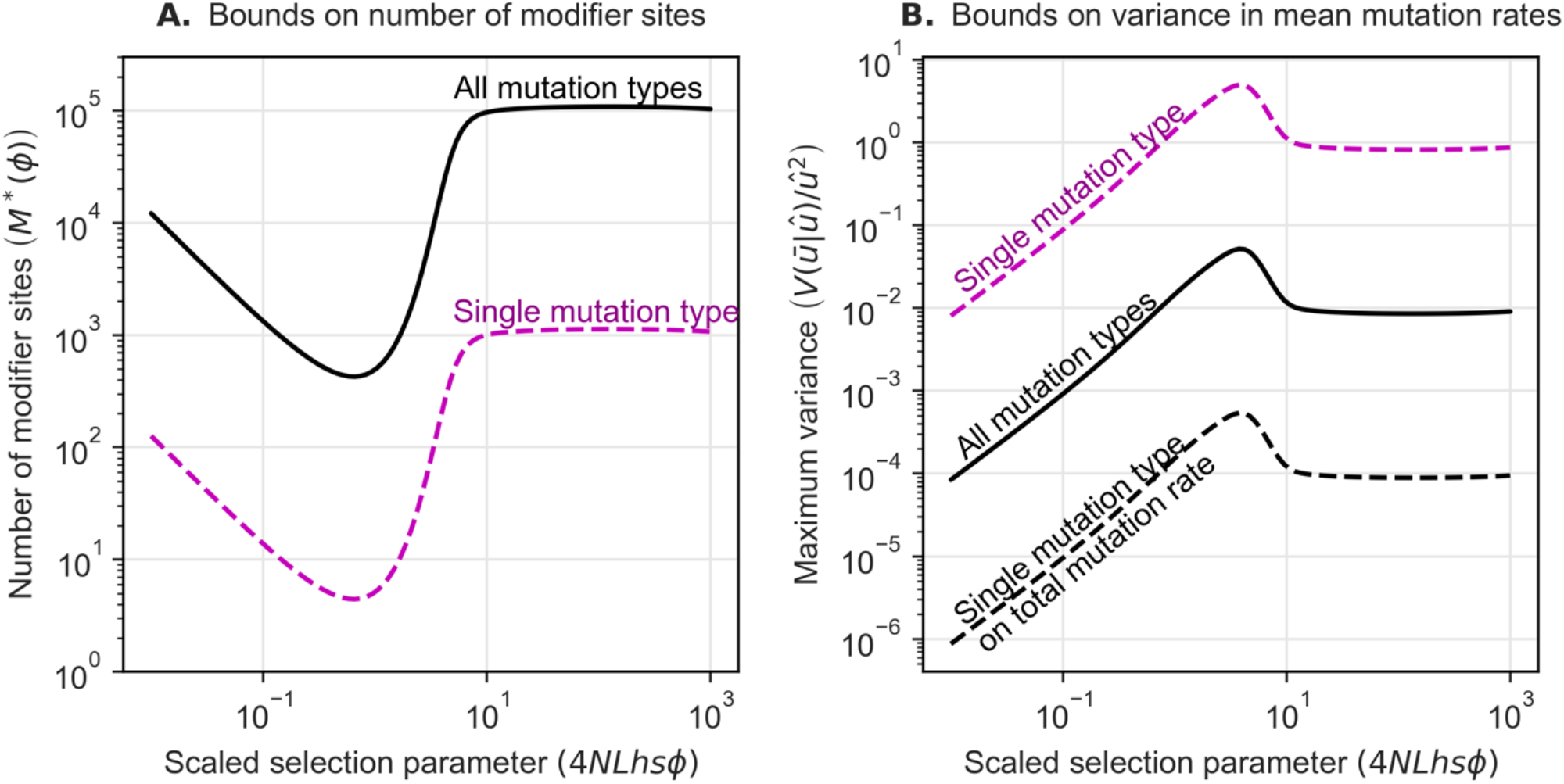
Upper bound on the number of modifier sites (A) and on variance in mean mutation rates (B), assuming that the expected mutation rate equals the estimated rate in humans. We show both quantities for modifier sites that affect one of 96 mutation types (dashed lines) and for those that affect all mutation types (solid lines). For modifier sites that only affect a single mutation type, we show both their effect on variance at sites they affect and on the total mutation rate. Each quantity was calculated using Eqs. 16 and 17. We assumed scaled parameter values roughly based on humans with *N* = 2 · 10^4^ (see *Parameters* and Table 1).

The variance in mean mutation rates conditional on a fixed expectation exhibits similar selection regimes as it does without this conditioning (Fig. 2). In the effectively neutral regime, selection negligibly affects the frequency of mutator alleles, so *V*(*q*)/*E*(*q*) remains approximately constant, and *V*(*ū*|*û*) increases linearly with the effect size, *ϕ* (Eq. 17). When mutators are strongly selected (i.e., under mutation-selection balance), then *E*(*q*)*∝*1/*ϕ* and *V*(*q*)*∝*1/*ϕ*^2^, as *s*^∗^ ∝ *ϕ*. Consequently, *V*(*ū*|*û*) is approximately constant. The conditional variance *V*(*ū*|*û*) is maximized in the nearly neutral regime. Using our estimates of human parameters (Table 1) and the upper bounds on the number of modifier sites (*M*^∗^(*ϕ*)), the maximum variance in the total mutation rate (relative to the expected rate squared) is ∼5%. However, if modifier sites only affect the rate of one of 96 mutation types, the maximum variance in that focal rate (again relative to the expected rate squared) is ∼500%. As expected, varying other parameters that make up the scaled selection parameter yields similar results (Fig. S2). We also find that this conditional variance behaves similarly when we allow mutator alleles to increase the mutation rate at both selected and modifier sites (Fig. S5).

### Interpreting the variance in mean mutation rate across populations

Our analytic results overestimate the variance expected among human populations. The variance that we calculate (Eqs. 14 and 17) reflects variation in mean mutation rates at a given point in time among independently-evolving populations at steady state. At steady state, most of the variance due to weakly selected modifier sites arises from differences in the number of fixed rather than segregating mutator alleles (because 4*Nμ* ≪ 1). However, the timescale of turnover of fixations at weakly-selected modifier sites is on the order of the expected time between neutral substitutions, 1/*μ* generations, which is much greater than the split time among human populations. Therefore, we expect few fixed differences at weakly selected modifier sites among human populations. In turn, strongly selected modifier sites never fix for the mutator allele and those that segregate turn over on a timescale proportional to the expected sojourn time, 1/*s*^∗^ generations (Fig. S3). Therefore, we expect strongly selected mutator alleles segregating in different human populations to be largely independent. However, observed mutation rate differences among extant populations typically reflect averages over many generations ancestral to the sample. Given the rapid turnover at strongly selected modifier sites, some of the difference in mean mutation rates may average out. For these reasons, we cannot rely on our analytic results to assess whether evolution at modifier sites can explain differences in mutation rates observed among human populations.

### Variation among human populations

We therefore rely on simulations to address this question. In the standard model that we implement, each modifier site affects the mutation rate at one of 96 mutation types and thus only affects 1/32 of selected sites. We constrain the number of modifier sites and baseline mutation rate, *u*_0_, such that the expected mutation rate (at steady state) at each mutation type is *û* = 1.25 · 10^−8^/3 per bp per generation (Eq. 16). We simulate 15 · 10^3^ mutator allele trajectories under the S&D demographic history for African and European populations (Fig. S4; see *Simulations* for details). For a given effect size *ϕ* and number of modifier sites *M*, we then resample these mutator trajectories to generate 10^5^ sets of 96 mean mutation rate trajectories (one for each mutation type), which include a shared part that occurs in the ancestral population, a part that occurs only in the European population, and a part that occurs only in the African population.

We use these sets of mutation rate trajectories to ask whether evolution at modifier sites could cause large, peak-like differences in mutation rates, such as the one observed for *TCC* → *TTC* mutations in European but not African populations. We reason that a peak would be reported regardless of the mutation type, population, and timing in which it occurred, and therefore examine sets of 96 trajectories jointly. We quantify the “peak pattern” using the two kinds of measures that were applied to human data in previous work.

Our first measure is based on the Harris and Pritchard (2017) enrichment statistic, which quantifies population differences in the proportion of segregating sites corresponding to a given mutation type. We mimic both their dataset and analysis as described in *Tests for variation in mutation rates between populations*. In brief, for each set of mutation rate trajectories, we generate realizations of mutation rate enrichments by dropping mutations on a simulated ancestral recombination graph (ARG) and recording the largest enrichment per set (see SI Section 4). We repeat this procedure for given values of *M* and *ϕ* to generate an empirical distribution of enrichment values, which we use to estimate the probability that the largest enrichment per set of trajectories exceeds 1.1 (whereas the greatest enrichment estimated by Harris and Pritchard (2017) was approximately 1.16).

Our second measure follows Speidel et al. (2019), who estimated the average mutation rate corresponding to 96 different mutation types in log-sized time-intervals along lineages leading to present-day African and European individuals. We do not mimic the Speidel et al. (2019) inference step, but instead we calculate the average mutation rate in 14 non-overlapping log-size intervals (7 per population) along our simulated mutation rate trajectories. We then classify a *set of mutation rate trajectories* as (Fig. 3D): *peak-like* if it includes at least one peak-like mutation rate trajectory but no trajectories with multiple elevations; *multi-elevated* if any trajectories have multiple elevations; or *not-peaked* if no trajectories have substantially increased mutation rates (see *Tests of variance between populations* for details). Finally, we estimate the probabilities that a set of mutation rate trajectories fits a given category for different values of *M* and *ϕ*.

**Figure 3.**
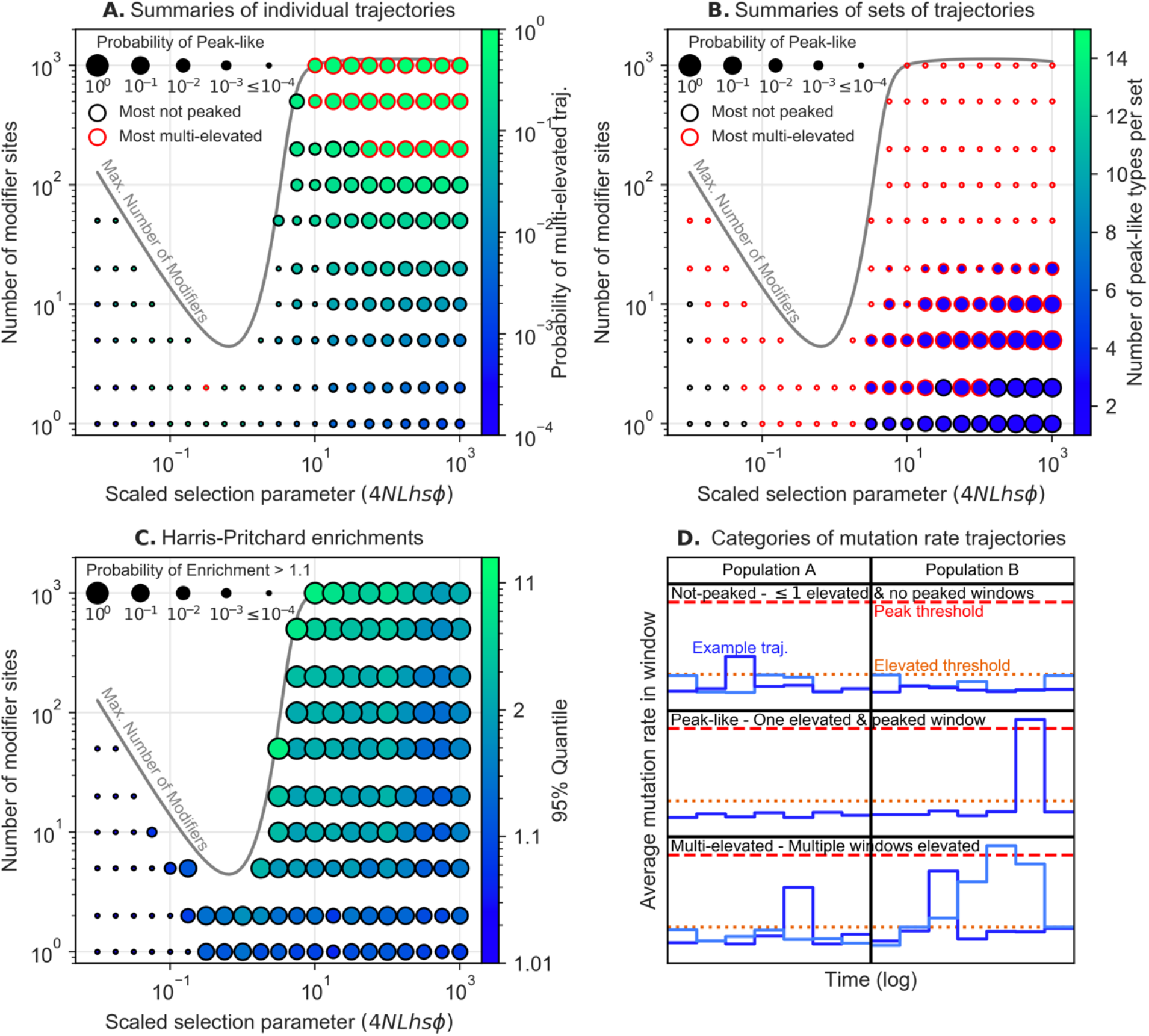
Summary of parameter ranges where modifier sites generate population specific peaks in mutation rates. For a given *M* and *ϕ*, we show the probability that (A) a single mutation rate trajectory or (B) sets of 96 trajectories are peak-like, or the probability that (C) the largest enrichment per set is greater than 1.1 (see *Tests of variance between populations*). (D) A cartoon illustrating how we categorize trajectories based on the number of windows that are above the elevated or peaked thresholds. For individual trajectories (A), we show the probability of being multi-elevated (interior color). For sets of trajectories (B), we show the expected number of peak-like trajectories in a peak-like set (interior color). For both (A and B), we show whether most trajectories or sets are not-peaked or multi-elevated (edge color). For enrichments (C), we show the 95% quantile of enrichment values (interior color). We vary *ϕ* to span the range of selection parameters and vary *M* between one and the maximum number of modifier sites possible, *M*^∗^(*ϕ*) (Eq. 16). In Figs. S11-14, we show similar results for models that relax some of our simplifying assumptions.

By these approaches, we find that for some parameter ranges, modifier sites frequently generate the differences observed among human populations (Fig. 3). Specifically, a few (*M* ≪ *M*^∗^) effectively neutral modifier sites (4*NLhsϕ* ≪ 1) cause only minor enrichments and almost no peak-like trajectories. However, the cumulative effects of many such modifier sites (*M*∼*M*^∗^) produces substantial differences in mutation rates between populations, which generate large enrichments, but such sites turn over too slowly to produce substantial differences in rates between populations that are contained to a single time interval. Thus, no number of effectively neutral modifier sites can produce peak-like trajectories. Modifier sites under moderate selection (1 ≤ 4*NLhsϕ* ≤ 10) turnover more rapidly and have greater effect sizes (Fig. S3), which allows them to produce larger enrichments and few peak-like trajectories. Modifier sites under stronger selection (4*NLhsϕ* > 10) turnover rapidly enough to allow for both large enrichments and frequent peak-like trajectories. Increasing the number of such modifier sites increases the frequency and magnitude of enrichments as well as the frequency of both peak-like and multi-elevated trajectories. Consequently, too many such modifier sites per mutation type (e.g., > 20) cause large enrichments but also almost always cause multi-elevated sets of trajectories. Putting these observations together, we conclude that a moderate number (≤ 10 per mutation type) under moderate to strong selection can generate large and localized increases in mutation rates that result in both large enrichments and peak-like sets of trajectories.

Evolution at modifier sites can also give rise to the observed enrichments and peaks under extensions of our basic model in which we consider: 1) mutation types with different expected rates (e.g., mutations at CpG sites whose rate is an order of magnitude greater than at other sites (Nachman and Crowell 2000; Kondrashov 2003; Kong et al. 2012)) (Fig. S12), 2) mutator alleles that are selected for their effects on somatic as well as germline mutation rates (Fig. S13); and 3) modifier sites that affect multiple mutation types (e.g., preferentially affecting 6 out of the 96 possible mutation types), which reflects the fact that different mutation types can arise from common underlying processes (Helleday et al. 2014; Alexandrov et al. 2020; Seplyarskiy et al. 2020) (Fig. S14). The parameter ranges with both large enrichments and peak-like sets of trajectories vary somewhat among models, and they vary noticeably with our criteria for peak-like trajectories (Fig. 4, see Fig. S11 for definitions of peak-like). When we use a more relaxed definition of peak-like trajectories (compared to the one used in Fig. 3), the parameter ranges expand but are still restricted to those under moderate to strong selection, which allow for sufficiently rapid turnover.

**Figure 4.**
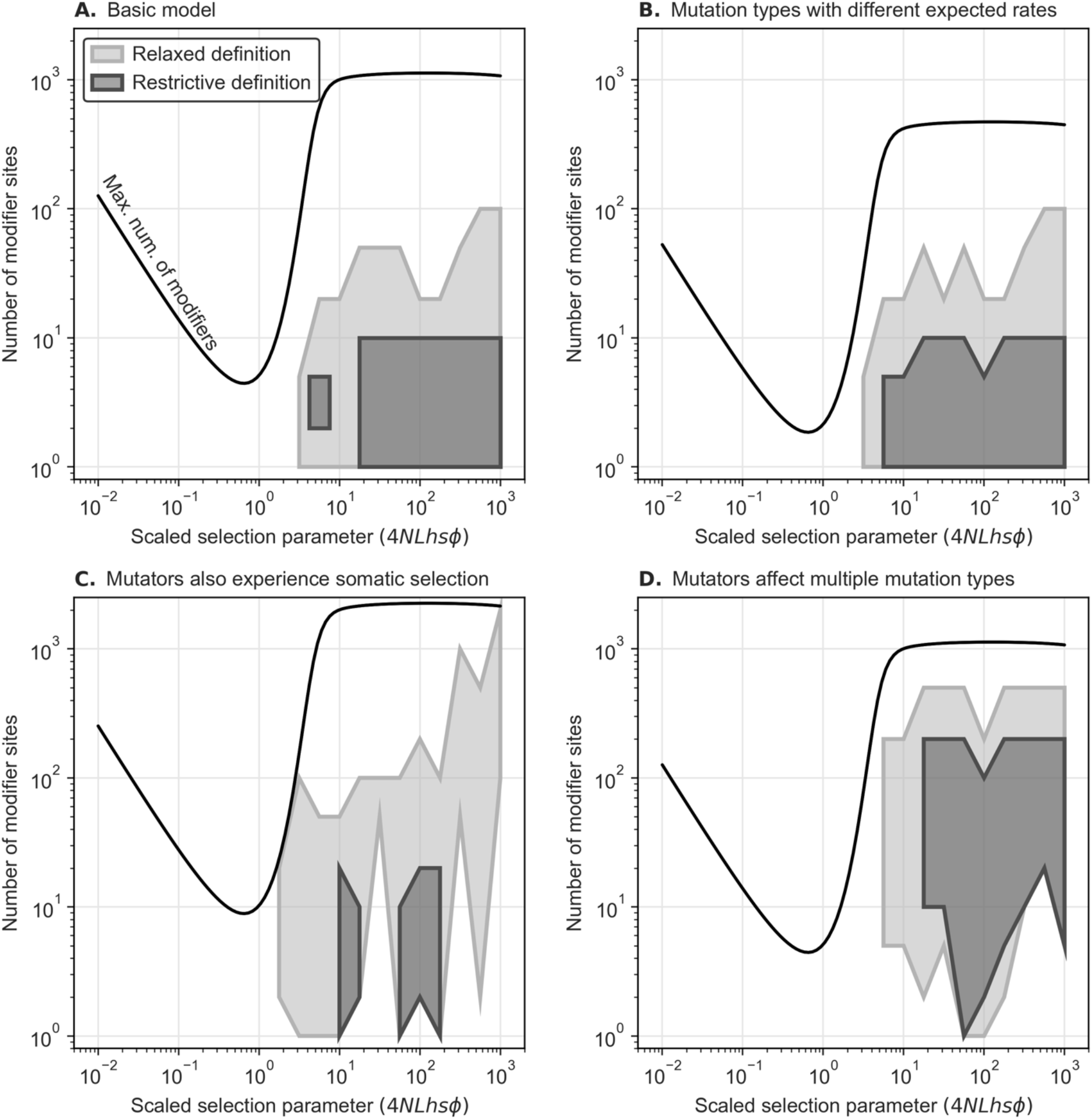
Parameter ranges in which both enrichments and peak-like sets occur with probability ≥ 0. 01 under different models and criteria for peak-like trajectories. (A) Our basic model. (B) A model in which mutation types have different expected rates. (C) A model in which mutator alleles are selected on due to their effects on germline and somatic mutation rates. (D) A model in which mutator alleles affect multiple mutation types. For all models, we show the parameter ranges for our restrictive (dark grey) and relaxed (light grey) definitions of peak-like trajectories (See *Tests of variance between populations* and Fig. S11 for definitions). See Figs. S12-S14 for additional details about and results for each model (akin to Fig. 3). We calculate the maximal number of modifier sites for a given scaled selection parameter (which varies across models) from Eq. 16. We calculated probabilities as we did for the basic model (see *Tests for variation in mutation rates between populations* and Fig. S11).

## Discussion

Our analyses suggest that the variation in mutation rates and spectra among human populations could have arisen predominantly from the evolution at modifiers (potentially in addition to changes in life history traits and environmental exposures; as proposed in e.g., Harris 2015; Mathieson and Reich 2017; Macià et al. 2021). This variation arises from the randomness of genetic drift and mutation at modifiers, which causes transient fluctuations in mutation rates and allows otherwise similar populations to evolve different mutation rates. Specifically, evolution at an intermediate number of modifier sites under moderate to strong selection (e.g., *M* ≤ 10 per type and 4*NLhsϕ* > 1) could have caused both the enrichments and peak-like increases in mutation rates observed between African and European populations (Harris 2015; Moorjani et al. 2016; Harris and Pritchard 2017; Mathieson and Reich 2017; Narasimhan et al. 2017; Aikens et al. 2019; Speidel et al. 2019; Goldberg and Harris 2019). Thus, our modeling recasts the question about the role of genetic modifiers in terms of their genetic architecture—i.e., about whether such parameter values are plausible.

We can potentially learn about the genetic architecture by studying mutation rate variation within populations. Unlike differences among populations, variation within a population arises solely from segregating modifier sites rather than substitutions. Specifically, the expected contribution of a single modifier site to variance in mutation rates is 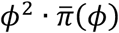 (Eq. 11), where *ϕ* is the effect size and 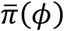 is the expected heterozygosity at the site. To understand how this contribution depends on modifier effect size, we consider simple approximations in the two selection extremes. For effectively neutral modifier sites, 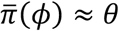, where *θ* = 4*Nμ* is the expected neutral heterozygosity; their expected contribution to variance therefore increases with *ϕ*^2^ and is approximately *ϕ*^2^ · *θ*. For strongly selected modifier sites, 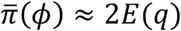, where *E*(*q*) ≈ *μ*/2*Lhsϕ* is the expected frequency of the mutator allele at mutation-selection balance. Their expected contribution to variance therefore increases linearly with their effect size 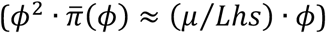, and the greatest contribution per site arises from large effect and thus rare segregating mutators (Fig. 5A). When we allow for the maximal number of modifier sites given a fixed expected mutation rate, *M*^∗^(*ϕ*), strongly selected modifier sites still contribute the most to variance within populations (Fig. 5B). This variance may allow us to identify the footprints of such strongly selected modifier sites, and thus we may learn about their number and effect size.

**Figure 5.**
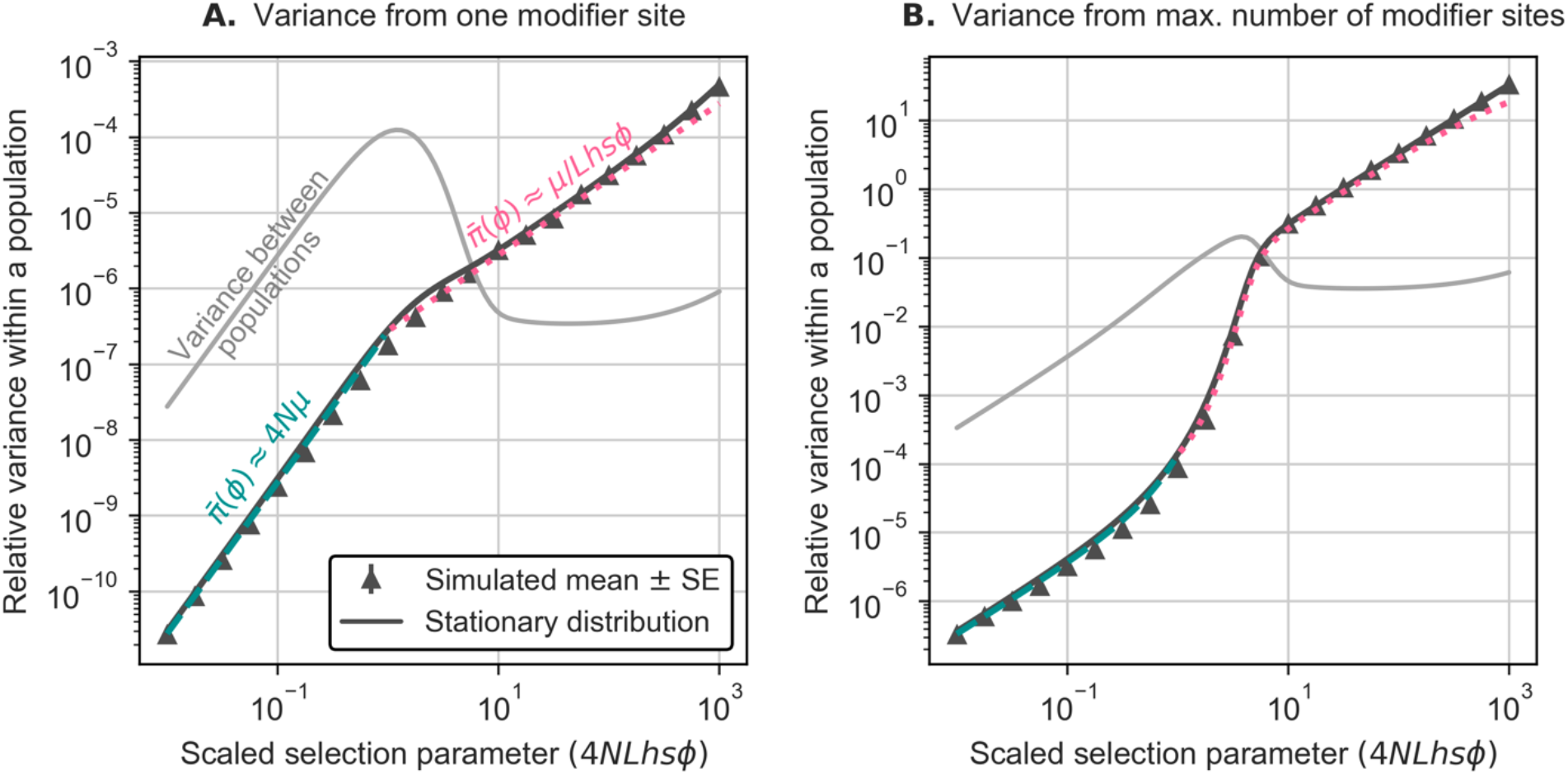
Variance in mutation rates within a population as a function of the effect size of modifier sites. We show the expected variance within a population (relative to a fixed expected rate, *E*(*V*(*u*|*q*))/*E*(*u*)^2^) at steady state arising from: (A) one modifier site and (B) the maximum number of modifier sites conditional on a fixed expected rate (Eq. 16). For simulated results with a single site (A), we estimate the average heterozygosity at modifier sites in simulations with *M* = 10^3^ (calculated as described in *Simulations*) and the contribution to variance within a population relying on Eq. 15. When assuming the maximum number of modifier sites (B), this estimated contribution and its SE are scaled by *M*^∗^(*ϕ*) (relying on Eq. 16 and assuming *û* = 1.25 · 10^−8^/*bp*/*gen*). In most cases, the SEs are too small to see. We calculate the analytical predictions from the expected heterozygosity, which is calculated by integrating over the stationary distribution, and relying on Eq. 15; we also show the results of the simple approximations described in the text. For comparison, we show the corresponding variance between populations, relying on Eq. 14. Both simulated and analytical results assume *N* = 10^3^ with other parameters chosen to match population-scaled values in humans (see *Parameters* and Table 1). These results are insensitive to plausible variation in model parameters (Fig. S2).

Seoighe and Scally (2017) suggested that mutator alleles segregating within a population could be identified based on the excess mutations linked to them. Closely linked sites are transmitted with the mutator allele and thus experience elevated mutation rates for more generations than loosely linked sites. However, a trade-off between the lifespan of a mutator allele and its effect size limits the expected number of mutations linked to mutator alleles such that this approach is likely underpowered (as we detail in SI Section 5).

An alternative approach may be to identify the footprints of strongly selected mutator alleles in pedigree studies of human mutation rates. We expect a parent that is heterozygous for a mutator allele with effect size *ϕ* to pass on *ϕδL*_*G*_ extra, de novo mutations to their children, where *L*_*G*_ is the number of sites (both selected and neutral) in the genome and *δ* is the proportion of sites within the genome affected by the mutator allele. If the mutator preferentially affects specific mutation types, it would also change the mutation spectrum. The effects of such a mutator could be detected if: 1) the excess number mutations that it causes is sufficiently large to be identified against the background of other sources of variation in the study, and 2) a parent carrying it is sampled in the study. For the purpose of illustration, we assume our basic model with a constant population size and plausible human parameter values (Table 1), that all de novo mutations are correctly identified, and that the number of de novo mutations in a proband is Poisson distributed with mean *L*_*G*_ (*u*_*M*_ + *u*_*P*_), where *u*_*M*_ and *u*_*P*_ are the maternal and paternal mutation rates. Thus, a proband with parents that carry a single mutator allele with effect size *ϕ* is expected to inherit *L*_*G*_ (2*u*_0_ + *δϕ*) de novo mutations. For illustration, we test for an increase in the number of de novo mutations in a trio under a simple null hypothesis, where there are no modifier sites (i.e., *M* = 0) and the number of de novo mutations is Poisson distributed with mean 2*L*_*G*_*u*_0_ (here *u*_0_ = *û*). Under these assumptions, we expect that a proband with parents that carry a single mutator with 4*NhsLϕ* ≥ 15 would have a significantly greater number of de novo mutations than expected (with *p* < 0.05), and that the significance level will increase with effect size (Fig. 6A). A single segregating mutator allele with such a large effect would be strongly selected against and rare. Specifically, given a sample of 1500 trios, on par with current human pedigree studies (Jónsson et al. 2017; Goldmann et al. 2018; Kessler et al. 2020; Rodriguez-Galindo et al. 2020), it is unlikely to be sampled (see SI Section 6). However, if the number of modifier sites is sufficiently large, then one or more mutators are likely to be sampled (Figs. 6B and S15, see SI Section 6 for more details). These considerations suggest that in principle, we should be able to identify the effects of mutators for a wide range of numbers (*M*) and effect sizes (*ϕ*) of modifier sites with current study designs (Fig. 6).

**Figure 6.**
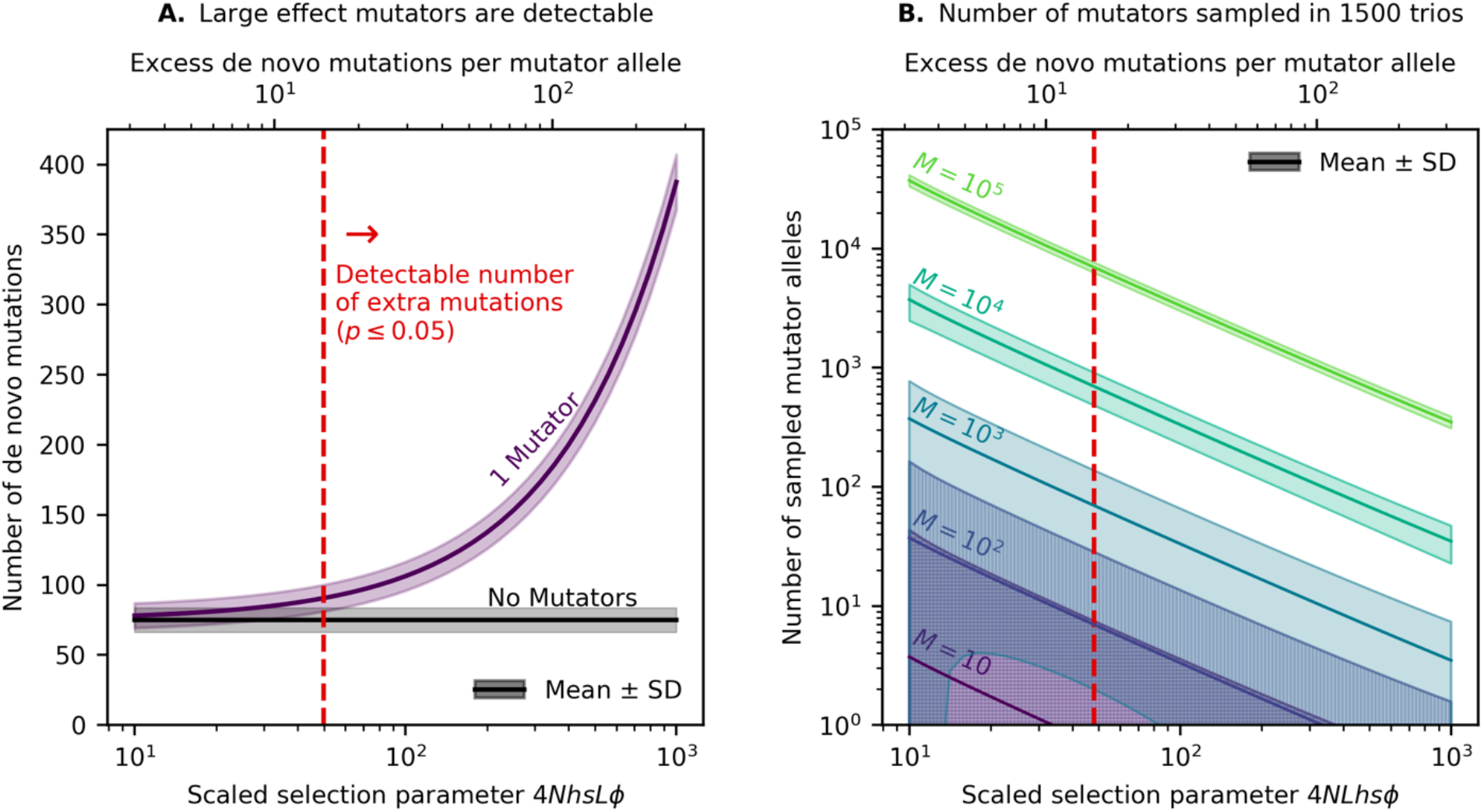
Identifying the effects of mutator alleles in a pedigree study of mutation rates. A) The expected number of de novo mutations (± 1 SD) in trios with and without a mutator. B) The expected number of mutators sampled (± 1 SD) in a study with 1,500 trios, based on our steady state approximations (Eq. 8 and SI Section 6). The vertical red lines denote the minimum effect size at which we expect a mutator to be detectable (with *p* < 0.05). For the purpose of this illustration, we assume a simple null model of mutations in the absence of mutators and ignore potential complications. We further assume that mutators affect all mutations (instead of, e.g., a specific mutation type) and rely on parameter values that are plausible for humans (Table 1) while varying *ϕ*.

We expect this qualitative conclusion to hold when we relax our simplifying assumptions. For example, while we rely on our steady state approximations, the expected number of sampled mutator alleles should be insensitive to recent population growth (Simons et al. 2014; Simons and Sella 2016). Other sources of mutation rate variation that we have ignored (e.g., parental age effects) could potentially be accounted for by a more realistic null model. Errors in identifying mutations in pedigree studies decrease the power to identify the effects of mutators, but we still expect them to be identifiable if our sample sizes or mutator effect sizes are sufficiently large. Thus, our calculations suggest that the footprints of strongly selected mutators will likely be detectable as pedigree study sample sizes increase allowing us to learn about mutator effect sizes and prevalence.

One of our main simplifying assumptions throughout was to ignore the selection on mutator alleles that arises from their effect somatic mutation rates. The form and strength of this selection should depend on 1) the effect of mutators on somatic mutation rates and 2) the effect of somatic mutations on fitness, both of which are poorly understood. Somatic and germline mutations arise from an interplay of damage, repair, and replication, which vary among cell lineages (Gao et al. 2016; García-Nieto et al. 2019; Alexandrov et al. 2020). Consequently, the effect of mutator alleles on somatic mutation rates are expected to vary across tissues and cell types (e.g., across neurons, colon crypts, and sperm). The fitness effects of somatic mutations may also be non-trivial. For example, the multi-hit model of carcinogenesis postulates that individual mutations in cancer driver genes may be tolerated whereas multiple mutations cause cancer (Knudson 1971; Knudson 2001; Anandakrishnan et al. 2019), which would suggest a non-linear relationship between somatic mutation rates and fitness (see, e.g., Lynch 2008). We can consider the potential effects of somatic selection on the evolution at modifier loci qualitatively. In the extreme case in which the somatic effects of mutator alleles are lethal, these alleles would have no chance to affect germline mutation rates. In the other extreme, in which selection is predominated by the effects in germline, we would largely recover our results without somatic selection. In between these extremes, the importance of somatic selection depends on its magnitude relative to germline selection, where this relative magnitude may vary with mutator effect size (see, e.g., Lynch 2008). In Fig. S14, we consider a simple model in which we assume that the strength of selection due to somatic and germline effects is similar. In that case and more generally, the addition of somatic selection reduces the frequency of mutator alleles and thus their impact on the expected mutation rate and the variance in rates within and between populations (Eqs. 13-15, Fig. S2). In turn, the reduced effect of individual modifier sites also allows for a greater number of modifier sites conditional on a given estimated mutation rate (Eq. 16), where the reduced effect and greater number of modifier sites may cancel out to still allow for modifier sites to explain the observed variation among populations and potentially to be identified in pedigree studies (Fig. S14). While these considerations suggest that our qualitative conclusions about variation in germline mutations within and among populations should hold, the potential effects of somatic selection on evolution at modifier loci merit their own study,

## Conclusion

Further extensions notwithstanding, our work lays the foundations for understanding of how genetic modifiers of germline mutation rates evolve across short evolutionary timescales and affect variation in mutation rates within and among populations. We derived closed forms for these effects for a population of constant size at steady state, which allowed us to characterize how they depend on basic evolutionary parameters. We investigated whether modifier sites can account for the properties of the mutation spectra observed in European and African populations, such as the observed peak in rates of *TCC* → *TTC* mutations in Europeans. Although we cannot rule out alternative hypotheses for the observed mutation rate variation, our findings suggest that a moderate number of modifier sites subject to moderate to strong selection could account for these patterns and that such modifier sites may soon be identified through sufficiently well-powered pedigree studies of mutation in humans.

## Acknowledgements

We thank Molly Przeworski for helpful discussions throughout this work. We also benefited greatly from discussion with Ipsita Agarwal, Peter Andolfatto, Laura Hayward, and Felix Wu and from comments on the manuscript from Molly Przeworski. This material is based upon work supported by the National Science Foundation Graduate Research Fellowship under Grant No. DGE1644869.

