## Supplemental Information for "The impact of genetic modifiers on variation in germline mutation rates within and among human populations"

### Table of contents

1. The excess number of mutations linked to a mutator allele ... 3
2. A correction for weakly selected mutators in finite populations ... 5
3. Stationary distribution ... 7
4. Simulating polymorphism datasets ... 9
5. Bounds on density of extra mutations linked to mutator alleles ... 11
6. Identifying mutator alleles in pedigree studies ... 11
   6.1 Expected number of mutators sampled. ... 12
   6.2 Variance in the number of mutators sampled ... 12
   6.3 Probability of having at least one mutator segregating in the sample. ... 12
7. References ... 13
8. Supplementary Figures ... 14

### List of supplementary figures

Fig. S1. Comparison of Kimura's and our approximations ... 14
Fig. S2. Sensitivity of our results to plausible variation in model parameters. ... 16
Fig. S3. Turnover rates at modifier sites.. ... 16
Fig. S4. Our model for the demographic history of European and African populations ... 18
Fig. S5. Variance in mean mutation rate when modifiers affect their own mutation rates. . 19

Fig. S6. Generating summaries of the ARG used to simulate polymorphism datasets. .......... 20
Fig. S7. Summaries of the simulated ARG as a function of time. .......... 21
Fig. S8. The Harris-Pritchard enrichment corresponding to a pulse in mutation rate .......... 22
Fig. S9. Sensitivity of expected mutation rates to changes in population size. .......... 23
Fig. S10. Sensitivity to weak dependencies among mutator trajectories. .......... 25
Fig. S11. Sensitivity to the definition of peak-like mutation rate trajectories .......... 26
Fig. S12. Sensitivity to heterogeneity in rates across mutation types. .......... 29
Fig. S13. Modifiers that affect the rates of several types of mutation. .......... 30
Fig. S14. A model with somatic selection against mutator alleles. .......... 33
Fig. S15. Probability of sampling a mutator allele in a pedigree study with 1500 trios .......... 34

### 1. The excess number of mutations linked to a mutator allele

Our aim is to calculate the excess number of deleterious mutations linked to mutator alleles in order to later approximate the average strength of selection acting against them. To this end, we extend Felsenstein's (Felsenstein 1974) classic derivation showing that at mutation-selection balance in an infinite population, the number of deleterious alleles that individuals carry follows a Poisson distribution with mean $2Lu/hs$. We assume mutator alleles are introduced into the population at a sufficiently low rate and selection against them is sufficiently strong that the proportion of individuals carrying a mutator allele satisfies $Q \ll 1$ at steady state. We divide the proportion of individuals into *age classes* based on the age of the mutator alleles, $t = 0, 1, \dots$, and derive recursions for the expected excess of deleterious alleles at birth, $X(t)$, and the number of individuals, $K(t)$, in each class.

First consider the generation in which the mutators are introduced, $t = 0$. Given that $Q \ll 1$, we approximate the proportion of individuals with new mutators by $2\mu$. We further assume that mutators of any age always appear in heterozygotes, with one mutator and one wildtype allele. Because we assume that mutators do not affect the mutation rate in the generation in which they appear, the number of deleterious alleles carried by individuals in age class $t = 0$ follows a Poisson distribution with mean $2Lu_0/hs$.

Now assume that the number of deleterious mutations in individuals of age class $t$ follows a Poisson distribution with mean $2Lu_0/hs + X(t)$. Then, in the following age class, $t + 1$, the fraction of individuals and distribution of deleterious mutations follow from changes that occur in a single generation: selection, mutation, recombination, mendelian segregation, and fertilization (Felsenstein 1974). After viability selection, the number of mutations is Poisson distributed with mean $(2Lu_0/hs + X(t)) \cdot (1 - hs)$. The number of de novo deleterious mutations is Poisson distributed with mean $2L(u_0 + \phi)$. After free recombination and Mendelian segregation, the number of deleterious alleles on the gamete carrying the mutator allele remains Poisson distributed with a mean that is halved. Lastly, the number of deleterious mutations on the wildtype gamete that is paired with the mutator gamete is Poisson distributed with mean $Lu_0/hs$. Thus, the excess number of mutations in age class $t+1$ is Poisson distributed with mean $X(t+1)$, where

$$2Lu_0/hs + X(t+1) = \frac{1}{2} \cdot \left[\left(2Lu_0/hs + X(t)\right) \cdot (1-hs) + 2L(u_0+\phi)\right] + Lu_0/hs.$$

This leads to the following, simple recursion

$$X(t+1) = \frac{1-hs}{2} \cdot X(t) + L\phi, \tag{S1}$$

which alongside the initial condition, $X(0) = 0$, is solved by the sum of a geometric series

$$X(t) = L\phi \cdot \sum_{i=0}^{t-1} \left(\frac{1-hs}{2}\right)^i = L\phi \cdot \frac{1-[(1-hs)/2]^t}{1-(1-hs)/2} \tag{S2}$$

for $t \geq 1$. As we detail in the main text, Kimura (1967) approximated the excess number of deleterious mutations by its asymptotic value, i.e., he assumed $X(t) \cong 2L\phi$ (neglecting further terms on the order of $hs$).

Next, consider the expected number of mutator alleles in age class $t$ that arise from a single mutational origin, $K(t)$. We can calculate this expectation using the recursion

$$K(t+1) = K(t) \cdot w(t), \tag{S3}$$

with initial condition $K(0) = 1$, where $w(t) = \overline{w}(t)/\overline{w}$ is the average fitness of individuals in age class $t$, $\overline{w}(t)$, relative to the population average, $\overline{w}$. The average fitness of a class follows from the number of deleterious mutations carried by individuals in it, which is Poisson distributed with mean $2Lu_0/hs$ for wildtype individuals and Poisson distributed with mean $2Lu_0/hs + X(t)$ for individuals in age class $t$. The average fitness of the population is well approximated by the average for wildtype individuals, $\overline{w} \cong \exp(-2Lu_0)$, because mutator alleles have a negligible effect on mean fitness. The average fitness of individuals in age class $t$ is $\overline{w}(t) = \exp(-(2Lu_0 + X(t) \cdot hs))$. Taken together, the relative fitness is $w(t) = \overline{w}(t)/\overline{w} = \exp(-X(t) \cdot hs)$, which implies that the expected numbers in different age classes satisfy the recursion

$$K(t+1) = K(t) \cdot e^{-hsX(t)}, \tag{S4}$$

with initial condition $K(0) = 1$. We use the recursions for $X(t)$ and $K(t)$ to approximate the expected integral over the number of mutator alleles that arise from a single mutational origin

$$K = \sum_{t=0}^{\infty} K(t), \tag{S5}$$

by calculating the sum of $K(t)$ up to some $T \gg 1/2Lhs\phi$.

The effective selection coefficient of mutator alleles, $s^*$ , defined as the one that generates the correct expected mutator allele frequency (summed over all age classes) at mutation-selection balance, is precisely the inverse of $K$, i.e.,

$$s^* = 1/K \tag{S6}$$

(Kimura 1967). When selection is extremely strong (e.g., $s^* \gtrsim 0.1$), our approximation for the effective selection coefficient is more accurate than the one used by Kimura (1967) and Lynch (2008) (Fig. S1), as such mutators rarely live long enough to near the asymptotic excess of deleterious mutations (see above and main text). When selection is not quite as strong ($s^* \ll 1$), long-lived alleles dominate K such that the two approximations are similar with $s^* \cong 2Lhs\phi$. Lastly, in finite populations, when selection is comparable to or weaker than genetic drift ($2Ns^* \cong 4NLhs\phi \lesssim 1$), both approximations become inaccurate. In this regime, the effect of the mutator allele on mean fitness or the existence of individuals homozygous for the mutator allele can no longer be neglected. We can nonetheless interpret $s^*$ as the fitness reduction of individuals heterozygous for the mutator allele relative to the mean fitness of an idealized population without mutator alleles. Under this interpretation, our approximation is accurate throughout the range of mutator effect sizes.

### 2. A correction for weakly selected mutators in finite populations

When the effects of selection on the frequency of mutator alleles are lesser or comparable to those of genetic drift ($2Ns^* \cong 4NLhs\phi \lesssim 1$), mutators can ascend to high frequencies. Consequently, when we calculate the effective selection coefficient, we cannot assume that individuals homozygous for mutator alleles are rare or that mutators have negligible effects on the population's mean fitness.

The case of weakly selected mutators therefore requires us to revise our derivations. We do so noting that:

- The number of de novo mutations on gametes of individuals heterozygous for the mutator is Poisson distributed with expectation $L(u_0 + \phi)$.

- The number of de novo mutations on gametes of individuals homozygous for the mutator is Poisson distributed with expectation $L(u_0 + 2\phi)$.
- The expected number of de novo mutations on a random gamete – without conditioning on the genotype at the modifier site – has expectation $L(u_0 + 2\phi q)$
- A mutator allele will be found in heterozygotes with probability $(1 - q)$ and in homozygotes with probability $q$. Consequently, the expected number of de novo mutations on gametes of individuals carrying the mutator allele is

$$(1 - q)(L\phi + Lu_0) + q(2L\phi + Lu_0) = L\big(u_0 + \phi(1 + q)\big), \tag{S7}$$

and the expected excess number of mutations on such gametes relative to a random gamete is

$$L\big(u_0 + \phi(1 + q)\big) - L(u_0 + 2\phi q) = L\phi(1 - q). \tag{S8}$$

Additionally, we assume that the number of de novo mutations on random gametes and on gametes of individuals carrying the mutator allele are also Poisson distributed (with the aforementioned expectations). These are approximations, as the distributions are in fact mixtures of Poisson distributions corresponding to different genotypes.

When weakly selected mutator alleles ascend to high frequencies, the expected number of excess deleterious mutations associated with them depends on their frequency. Consider the expected number of excess deleterious mutations associated with a mutator alleles in age class $t$, $X(t)$. In this case, the number of excess mutations added each generation is $L\phi(1 - q)$ rather than $L\phi$ (Eq. S8). Namely,

$$X(t + 1) = X(t)\frac{1 - hs}{2} + L\phi\big(1 - q(t)\big) \tag{S9}$$

with initial condition $X(0) = 0$. We assume that the number of excess mutations $X(t)$ equilibrates much faster than the mutator's frequency $q(t)$ changes. This assumption should hold when $q(t)$ is substantial, because in this case $q(t)$ changes on the timescale of genetic drift whereas $X(t)$ equilibrates on the same timescale as the decay of transient linkage between mutator alleles and any excess mutations (which is very fast when we assume all sites can freely recombine). In turn, when $q(t)$ is very small, the dependence of $X(t + 1)$ on $q(t)$ is negligible as $1 - q(t) \approx 1$. Under this assumption, Eq. S9 becomes

$$X(t+1) = X(t)\frac{1-hs}{2} + L\phi(1-q) \tag{S10}$$

with asymptotic value, $X(\infty) \cong 2L\phi(1-q)$. By analogy with the derivation in the previous section and given the separation of time scales argument, we approximate the expected excess of deleterious alleles in any age class by its asymptotic value. Moreover, we approximate the relative fitness of a mutator allele with frequency $q$ by

$$w(q) = \bar{w}_M(q)/\bar{w}(q) \cong \exp\left(-2Lhs\left(u_0 + \phi(1+q)\right)\right)/\exp\left(-2Lhs(u_0 + 2\phi q)\right)$$

$$= \exp\left(-2Lhs\phi(1-q)\right),$$

where $\bar{w}_M(q)$ is the average fitness of individuals carrying the mutator allele, and $\bar{w}(q)$ is the average fitness in the population. The effective selection coefficient associated with a mutator allele with frequency $q$ is then

$$s^{**} = 1 - w(q) \cong 1 - \exp\left(-2Lhs\phi(1-q)\right) \cong 2Lhs\phi(1-q) \cong s^*(1-q) \tag{S11}$$

where $s^*$ is the approximate selection coefficient we derived for strongly selected mutator alleles in SI Section 1.

We use $s^*(1-q)$ as our approximation for the effective selection coefficient throughout the entire range of mutator effect sizes. When selection is weak, $s^* \cong 2Lhs\phi$ and thus $s^{**} \cong s^*(1-q)$; when selection is strong, $q \ll 1$ and thus $s^{**} \cong s^*$. Our approximation is similar to Kimura's for intermediate selection coefficients (when $2Ns^* \gg 1$ and $s^* \ll 1$) but produces substantially weaker and more accurate selection in the two extremes (Fig S1).

### 3. Stationary distribution

We now derive the diffusion approximation for the stationary frequency distribution at a modifier site based on the first two moments of change in mutator allele frequency in a single generation (Eqs. 6 and 7),

$$E(q) = -q(1-q)(1-q)s^* + \mu(1-2q) \text{ and } V(q) = \tfrac{q(1-q)}{2N}. \tag{S12}$$

In general, the distribution is approximated by

$$\Phi(q) = C\exp\left(2\int_0^q \frac{E(x)}{V(x)}dx\right)/V(q), \tag{S13}$$

where $C$ is a normalizing constant defined such that $\int_0^1 \Phi(q)\, dq = 1$ (Fisher 1923; Wright 1945; Kimura 1955; Ewens 2004). In our case, this takes the form

$$\Phi(q) = C\big(q(1-q)\big)^{4N\mu-1} e^{-4Ns^{*}q(1-q/2)}, \tag{S14}$$

which is an application of Wright's formula (Wright 1945). By integrating over this distribution, we can calculate any summary of mutator allele frequencies, including their effect on the mean and variance of the mutation rate.

However, when $4N\mu < 1$, which is the biologically relevant parameter regime, $\Phi(q)$ is singular at the boundaries $q = 0$ and 1, which causes the normalization constant $C$ to diverge. This problem arises from using a continuous approximation for discrete frequencies and is resolved by moving back to a discrete Markovian process to calculate the probability mass at the boundaries, $\Phi(0)$ and $\Phi(1)$ (Ewens 2004). Specifically, we calculate the probability at $q = 0$ by summing over the probabilities of being at state $i$, corresponding to frequency $i/2N$, $\Psi(i/2N)$, multiplied by the transition probability from state $i$ to 0 in the next generation, $\Pr(i \to 0)$, such that:

$$\Psi(0) \cong C\frac{\sum_{i=1}^{2N-1} \Psi(i/2N)\Pr(i \to 0)}{1 - \Pr(0 \to 0)}, \tag{S15}$$

where we neglect the transition probability from frequency 1 to 0. We approximate the probability of being at state $i = 1, \dots, 2N-1$ based on the continuous approximation, i.e.,

$$\Psi(i/2N) = C\int_{i/2N-1/4N}^{i/2N+1/4N} \Phi(q)\, dq. \tag{S16}$$

We first solve Eq. S16 in terms $\widetilde{\Psi}(i/2N) = \Psi(i/2N)/C$, with $i = 0, \dots, 2N-1$, such that it does not include the normalization constant $C$, and later solve for $C$.

We approximate the transition probability from $i$ to 0 as the transition probability from $i$ to 0 via Wright-Fisher sampling, $\Pr_{WF}(i \to 0)$, multiplied by the probability of having no mutation from the wild-type to the mutator state, $\Pr_M(0 \to 0)$. We neglect the probability of transitioning from $i$ to $j \neq 0$ via Wright-Fisher sampling and transitioning from $j \neq 0$ to 0 via mutation. The probability of having $j$ mutator alleles in the next generation under Wright-Fisher sampling follows the Binomial process:

$$j \sim Bin\left(2N, \exp\left(-s^*\left(1-\frac{i}{2N}\right)\right)\left(\frac{i}{2N}\right)\right), \tag{S17}$$

and specifically:

$$\Pr_{WF}(i \to 0) = \left(1 - \exp\left(-s^*\left(1-\frac{i}{2N}\right)\right)\left(\frac{i}{2N}\right)\right)^{2N}.$$

The probability of having $j$ mutators arise as de novo mutations given that 0 were sampled follows a Poisson process with mean $2N\mu$ and in particular:

$$\Pr_M(0 \to 0) = \exp(-2N\mu).$$

Thus, we find that:

$$\Pr(i \to 0) \cong \Pr_{WF}(i \to 0) \cdot \Pr_M(0 \to 0) \tag{S18}$$

$$= \left(1 - \exp\left(-s^*\left(1-\frac{i}{2N}\right)\right)\left(\frac{i}{2N}\right)\right)^{2N} \cdot \exp(-2N\mu).$$

We solve for $\tilde{\Psi}(0) = \Psi(0)/C$ numerically by substituting the expressions for $\tilde{\Psi}(i/2N)$ and $\Pr(i \to 0)$ (Eqs. S16 and S18 respectively) into Eq. S15; we solve for $\tilde{\Psi}(1)$ similarly. We then solve for the normalization constant, $C$, by requiring that

$$C\left(\tilde{\Psi}(0) + \tilde{\Psi}(1) + \int_{1/4N}^{1-1/4N} \Phi(q)\, dq\right) = 1.$$

A python script for calculating the stationary distribution and summaries of mutator allele frequencies is available for download at https://github.com/sellalab/Mutator.

### 4. Simulating polymorphism datasets

Here, we describe how we simulate the polymorphism datasets, which we then use to calculate the enrichment statistics (see T*ests for variation in mutation rates between populations*). We use msprime (Kelleher et al. 2016) to simulate the ancestral recombination graph (ARG), assuming a demographic model for Europeans and Africans based on Schiffels and Durbin (2014) (Fig. S4), a sample size of 500 individuals in each population (i.e., 1000 haploid genomes), a genome size of $3 \times 10^9$ base pairs (bp) divided into 100 equally sized chromosomes, and a sex-averaged recombination rate of $2 \cdot 10^{-8}$ per bp per generation. Msprime generates ARGs in continuous generations, so we round the time of each coalescence event to the nearest integer in order to obtain an ARG that is defined in discrete generations. We define the following summaries of the simulated ARG, corresponding to the number of opportunities for mutation at generation $t$ that would result in different kinds of polymorphisms within the sample (Fig. S6):

- $C(t)$ – The number of lineages at time $t$ that are private to Europeans (i.e., whose descendants in the sample are all European) summed over all marginal trees weighted by the number of base pairs each tree spans.
- $Y(t)$ – defined as $C$, but private to Africans.
- $A(t)$ – defined similarly to $C$ and $Y$, but for lineages that are ancestral to samples from both populations, not including lineages that ancestral to the entire sample (i.e., above the most recent common ancestor (MRCA) of the all 1000 individuals at a locus).

Given these summaries and a mutation rate trajectory corresponding to a given mutation type, $u(t)$, we choose the number of polymorphisms of this type in the European sample, $Z$, based on the distribution

$$Z(\text{European}) \sim Pois\left(\sum_{t=1}^{TMRCA} u_E(t)(C(t) + A(t))/32\right), \quad \text{(S19)}$$

where $u_E$ denotes the mutation rate trajectory describing mean mutation rates in the ancestral population and in the European population after the populations split and the division by 32 reflects our assumption of an equal number of each type of ancestral triplet (e.g., $TCC$). At most loci, the time to the MRCA is greater than the time at which our simulated mutation rate trajectories begin, so we assume that the mutation rate beforehand is the average over the given mutation rate trajectory. We choose the number of polymorphisms in the African sample based on a similar expression (replacing $C$ with $Y$ and $u_E$ with $u_A$, the equivalent mutation rate trajectory for African lineages in Eq. S19), and repeat this process for each mutation rate trajectory in a simulated set (see Simulations section). From each set of mutation rate trajectories, we obtain a single, corresponding dataset of polymorphisms. We use the same genome-wide simulated ARG to generate all of our polymorphism datasets, but in Fig. S7 we show that variation in the genome-wide ARG has a negligible effect on our results.

### 5. Bounds on density of extra mutations linked to mutator alleles

Seoighe and Scally (2017) suggested mutator alleles segregating within population could be identified by leveraging the excess mutations linked to them. A segregating mutator allele with effect size $\phi$ that arose $g$ generations ago would carry on average to $\phi g$ extra, derived (neutral) alleles per bp across the $2/(rg)$ bp haplotype that remained linked to it, where $r$ is the recombination rate per bp per generation (Seoighe and Scally 2017). For such a signature to be identifiable, the excess density of derived alleles has to rise above the level of background genetic diversity. This density is constrained by the trade-off between mutator lifespan and effect size. Strongly selected mutators persist for $g \approx 1/s^* \approx 1/(2hsL\phi)$ generations on average, at which point the expected excess density is $\phi g \approx 1/(2hsL)$. Given plausible human parameter values, this calculation suggests an upper bound of approximately 1 extra derived allele every 0.24Mb—three orders of magnitude smaller than average neutral heterozygosity (with $\bar{\pi}_0 \approx 0.001$ (Li and Durbin 2011)), so likely much too weak to be identifiable. In turn, weakly selected mutators should cause at most 1 extra derived allele every 0.012Mb (an order of magnitude below the average neutral heterozygosity), assuming conservative limits of $g \leq 4N$ and $\phi < 10/(4NLhs)$. These bounds would be lower if mutators face additional selection due to pleiotropic selection, selection against linked mutations, or selection against increased somatic mutation rates. Furthermore, we expect these results to be insensitive to recent population growth (Simons et al. 2014; Simons and Sella 2016). Thus, the theory derived here suggests this approach is likely underpowered across the entire parameter space, a result that should be insensitive to our specific assumptions.

### 6. Identifying mutator alleles in pedigree studies

Here, we calculate several quantities that describe the effects of mutator alleles in pedigree studies. To this end, we assume a population of constant size at steady state, and that modifier sites evolve independently.

**6.1 Expected number of mutators sampled.** First, we consider the number of mutator alleles sampled at a single modifier site. Given mutator allele frequency $q$, the number of mutator alleles sampled in $2n$ parents is distributed as:

$$Y_1 \sim Bin(4n, q). \qquad \text{(S20)}$$

In turn, the mutator allele frequency, $q$, follows the stationary distribution that we approximated in SI Section 3. Therefore, we expect to sample $4nE(q)$ mutator alleles at a single modifier site and $4nME(q)$ mutator alleles at $M$ modifier sites. We can only detect segregating alleles, so this calculation only holds for strongly selected modifier sites for which the probability of a mutator allele fixing in the sample is negligible.

**6.2 Variance in the number of mutators sampled.** Starting with a single modifier site, we apply the law of total variance to find that

$$V(Y_1) = V\big(E(Y_1|q)\big) + E\big(V(Y_1|q)\big) = (4n)^2 V(q) + 4n\big(E(q) - E(q^2)\big), \qquad \text{(S21)}$$

where $V\big(E(Y_1|q)\big)$ reflects variance from variation in mutator frequency, and $E\big(V(Y_1|q)\big)$ reflects variance from binomial sampling. Assuming independence among modifier sites, we find that

$$V(Y_M) = MV(Y_1) = M\left((4n)^2 V(q) + 4n\big(E(q) - E(q^2)\big)\right) \cong M(4n)^2 V(q), \qquad \text{(S22)}$$

where variance from mutator frequency dominates variance from binomial sampling. In Fig. 6B, we plot the corresponding standard deviation around the expectation to indicate the plausible range on the number of mutators sampled, i.e.,

$$4nME(q) \pm \sqrt{M\left((4n)^2 V(q) + 4n\big(E(q) - E(q^2)\big)\right)}. \qquad \text{(S23)}$$

**6.3 Probability of having at least one mutator segregating in the sample.** In Fig. S15, we show the probability of having at least one mutator segregating in the sample for different values of the scaled selection parameter and number of modifier sites. Given the stationary distribution of mutator frequency, we calculate this probability as the complement of having no mutator segregating in the sample, i.e.,

$$1 - \left(\int_0^1 \left((1-q)^{4n} + \mathrm{q}^{4\mathrm{n}}\right) \Psi(q) dq\right)^M. \qquad \text{(S24)}$$

### 8. Supplementary Figures

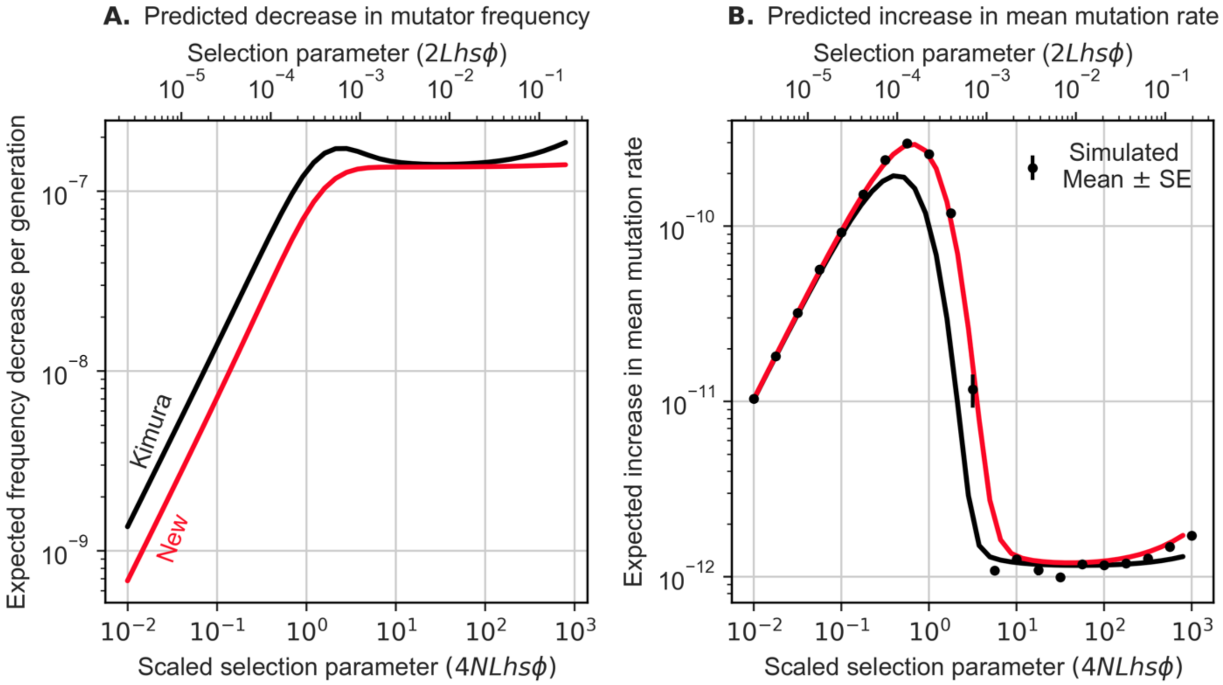


**Figure S1. Comparison of Kimura's and our approximations** for the effective selection coefficient of mutator alleles. (A) Expected change in mutator allele frequency per generation as a function of the scaled selection parameter. In Kimura's approximation $E(\Delta q_s) \approx e^{(1-2hsL\phi)}E\big(q(1-q)\big)$ and in our approximation $E(\Delta q_s) \approx -s^* E(q(1-q)^2)$. (B) Expected increase in mean mutation rates from a single modifier site as a function of the scaled selection parameter. Simulation-based estimates and SEs are calculated as described in *Simulations*. In both graphs, we assume a constant population size of $N = 1000$ with other parameters chosen to match population-scaled values in humans (see *Model Parameters* and Table 1). Because selection is weaker in our approximation than in Kimura's, we predict smaller changes in frequency due to selection and greater increases in mutation rates than Kimura did. Our approximation more accurately predicts the results of simulations, although the difference is negligible when mutators are effectively neutral ($4NLhs\phi \ll 1$) or when the two approximations are similar (e.g., $10 \leq 4NLhs\phi \leq 100$).

290

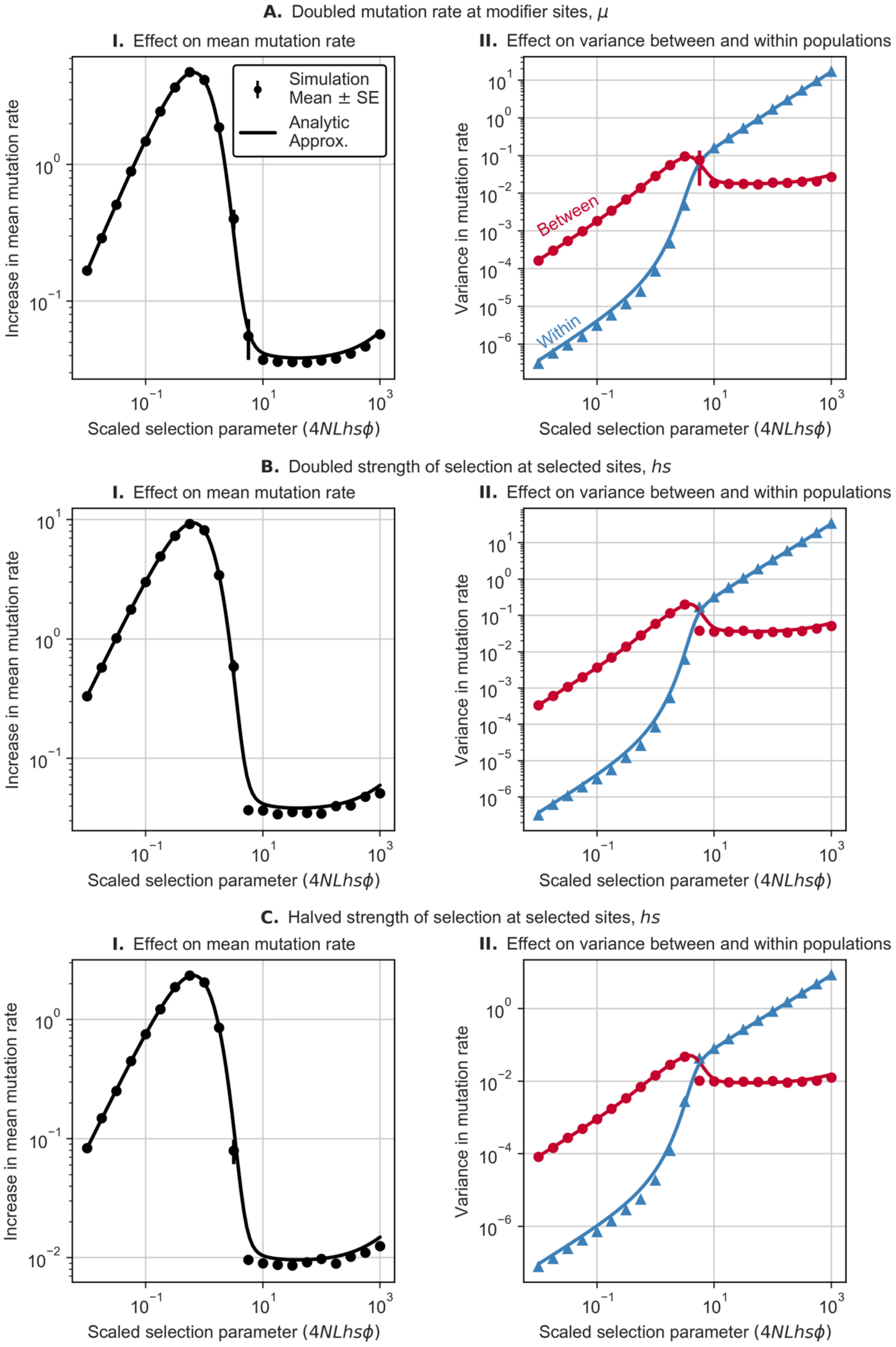

A. Doubled mutation rate at modifier sites, μ
I. Effect on mean mutation rate
Simulation
Mean ± SE
Analytic
Approx.
Increase in mean mutation rate
Scaled selection parameter (4NLhsϕ)
II. Effect on variance between and within populations
Variance in mutation rate
Between
Within
Scaled selection parameter (4NLhsϕ)
B. Doubled strength of selection at selected sites, hs
I. Effect on mean mutation rate
Increase in mean mutation rate
Scaled selection parameter (4NLhsϕ)
II. Effect on variance between and within populations
Variance in mutation rate
Scaled selection parameter (4NLhsϕ)
C. Halved strength of selection at selected sites, hs
I. Effect on mean mutation rate
Increase in mean mutation rate
Scaled selection parameter (4NLhsϕ)
II. Effect on variance between and within populations
Variance in mutation rate
Scaled selection parameter (4NLhsϕ)

**Figure S2. Sensitivity of our results to plausible variation in model parameters.** Similar to Figs. 1 and 6A, we show the effect of modifier sites on the mutation rate mean (left, I) and variance (right, II), as a function of the scaled selection parameter. We (A) double the mutation rate at modifier sites, $\mu$; (B) double the selection coefficient at selected sites, $\overline{hs}$; or (C) halve the selection coefficient. For all panels, we assume $M = 10^3$ modifier sites and use the same parameters as in Fig. 1, except for the one parameter we vary. Simulated quantities are calculated as described in *Simulations* using only 10 replicates per effect size*.* In most cases, the SEs are too small to see and hence are not shown.

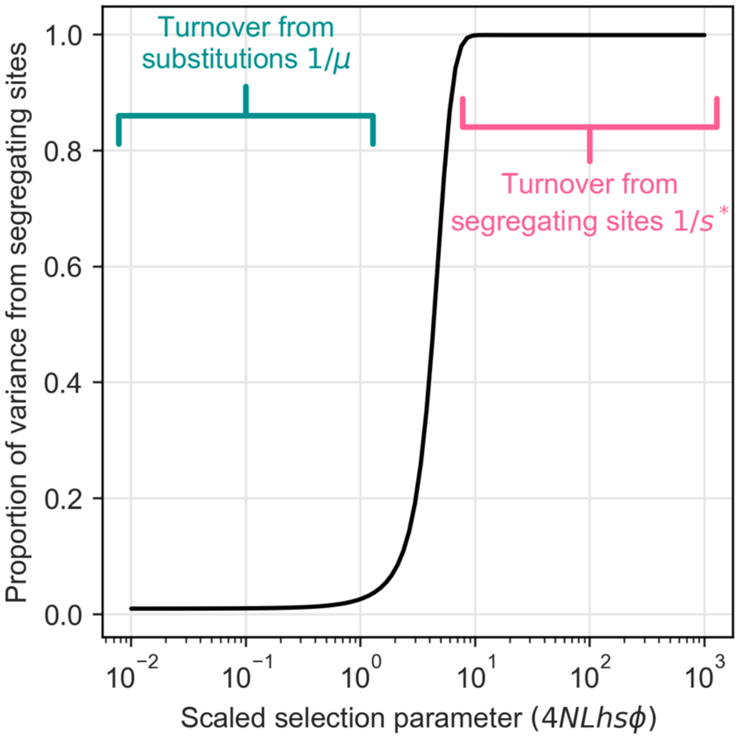


**Figure S3. Turnover rates at modifier sites.** We calculate the proportion of variance arising from segregating sites using our steady state approximations (Eq. 8), i.e., $V(q|0 < q < 1)/V(q) \cong \int_{1/4N}^{1-1/4N} \left(p - E(p)\right)^2 \Psi(p)\, dp / V(q)$. When modifier sites are effectively neutral ($4NLhs\phi \ll 1$), most of the variance in mean mutation rate arises from substitutions, which occur on a molecular evolutionary timescale of $\sim 1/\mu$ generations. When modifier sites are strongly selected ($4NLhs\phi \gg 1$), most of the variance arises from segregating sites, which turnover on a timescale of $1/s^*$ generations.

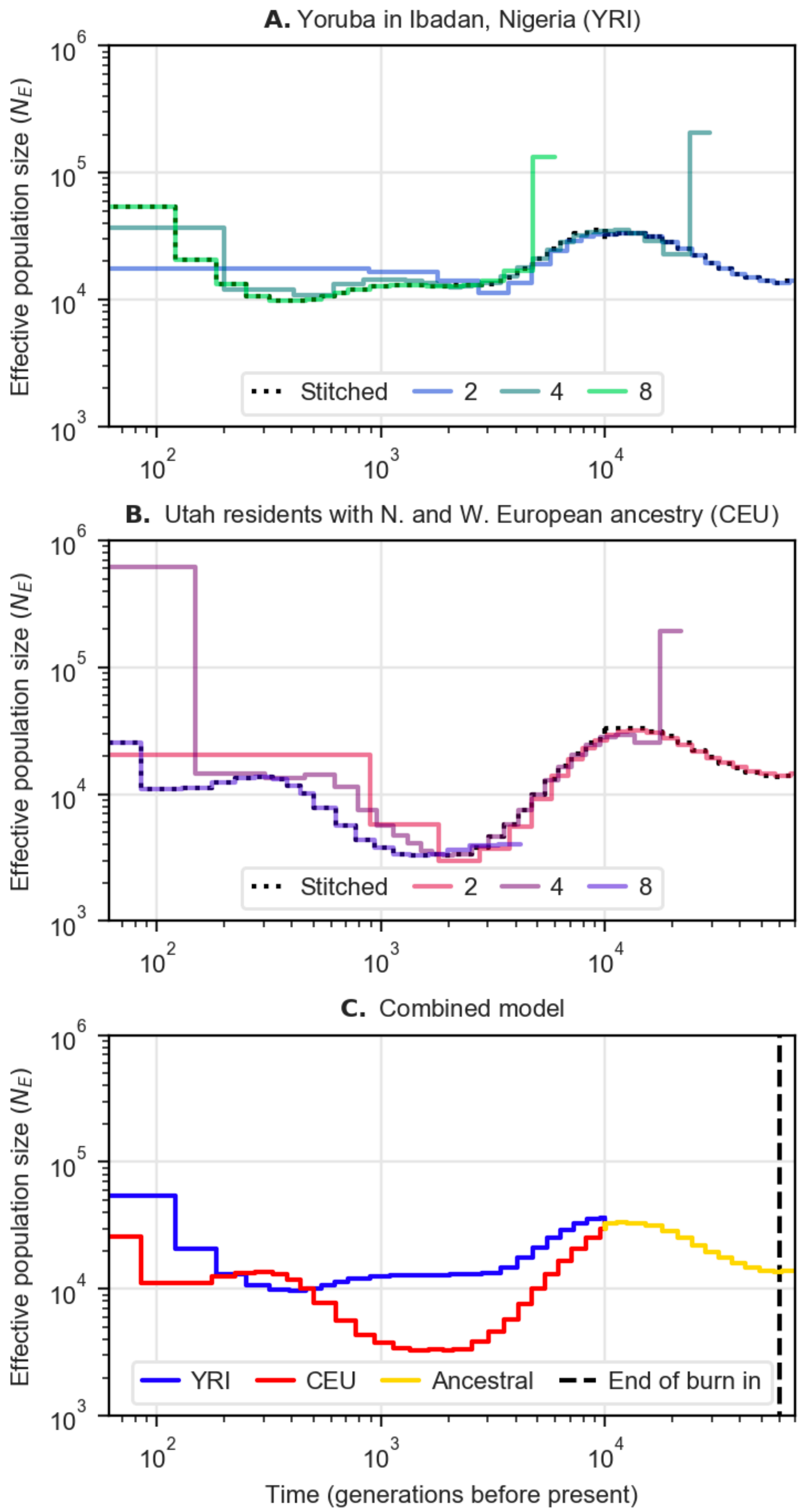
A. Yoruba in Ibadan, Nigeria (YRI)
Effective population size ($N_E$)
Stitched
2
4
8
B. Utah residents with N. and W. European ancestry (CEU)
C. Combined model
YRI
CEU
Ancestral
End of burn in
Time (generations before present)

307

**Figure S4. Our model for the demographic history of European and African populations.** We rely on the models inferred by Schiffels and Durbin for (A) Yoruba in Ibadan, Nigeria (YRI) and (B) Utah residents with Northern and Western European ancestry (CEU) (Schiffels and Durbin 2014). For each population, we show the past effective population sizes they inferred based on samples of 2, 4, or 8 haplotypes from the respective population. Using larger sample sizes improves the estimates in the recent past but biases estimates in the more distant past (Schiffels and Durbin 2014). We therefore visually 'stitch' the estimates based on different sample sizes together, as depicted by the black dotted curves shown in (A) and (B). We also estimate the split time between populations visually, based on the time at which the estimated population sizes in YRI and CEU begin to (approximately) overlap, and use the estimates in CEU for the ancestral population size prior to that time. Schiffels and Durbin report their inferences in years assuming a generation time of 20 years. Our simulations require discrete generations, so we convert back to generations assuming a generation time of 25 years and round to the nearest integer. (C) We show the resulting demographic model that we use, which we refer to as the S&D demographic history.

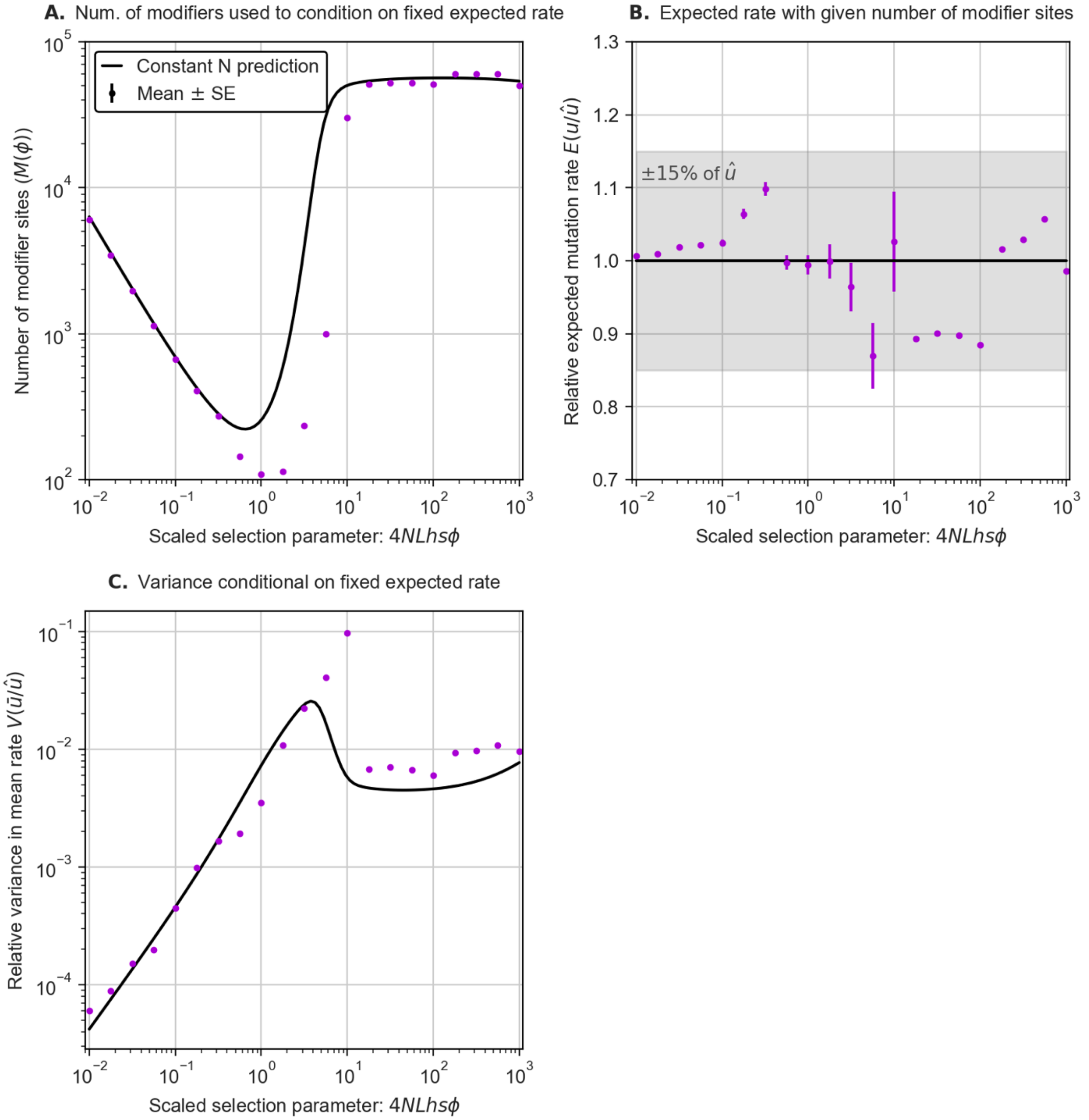


**Figure S5. Variance in mean mutation rate when modifiers affect their own mutation rates.** We consider a model in which mutator alleles increase the mutation rate at both selected and modifier sites. Otherwise, we assume our standard model with a constant population size $N = 1000$ (see *Model*), population scaled selection parameters corresponding to humans in the case where modifier sites have a constant mutation rate (Table 1), a baseline mutation rate of $u_0 = \hat{u}/2 = 1.25 \cdot 10^{-7}/2$ per bp per generation, and we set the number of modifier sites with effect size $\phi$, $M(\phi)$, such that the expected mutation rate approximately equals $\hat{u}$. Here, we cannot assume that modifier sites evolve independently of each other, so we use simulations to determine $M(\phi)$. Specifically, for each value of $\phi$, we run simulations with different numbers of modifier sites, $M$, estimate the expectation mutation rate $E(u|\phi, M)$, and keep adjusting $M$ until $E(u|\phi, M)$ is within 15% of $\hat{u}$. The number of modifier sites used in simulations, $M(\phi)$, and the corresponding estimated expected mutation rate as a function of the scaled selection parameter are shown in (A) and (B), respectively. In (C), we use simulations with $M(\phi)$ modifier sites to estimate the variance in mean mutation rate $V(\bar{u}|\phi, M(\phi))$. We calculate the estimated quantities and their SEs as described in *Simulations*; SEs are often too small to be seen. After conditioning on $E(u|\phi, M) \approx \hat{u}$, $V(\bar{u}|\phi, M(\phi))$ displays the same selection regimes that we saw with our standard model (Fig. 1). However, when selection is moderate ($4NLhs\phi \sim 10$), the variance is substantially greater than our analytic prediction. This may reflect transient surges in mutation rate in which segregating mutators amplify the rate at which other mutators arise, thus increasing the variance in mean mutation rate. This effect should be much weaker if individual modifier sites only affect the rates of a subset of mutation types, as the opportunity for such non-linear interactions among modifier sites would be greatly reduced.

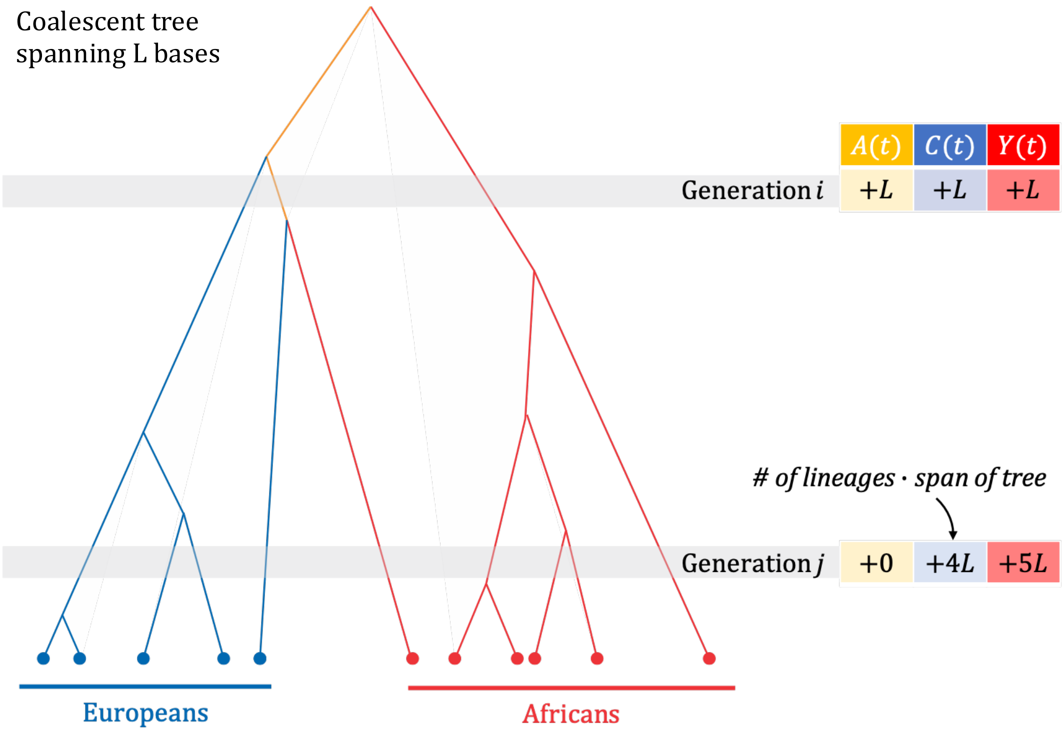


**Figure S6. Generating summaries of the ARG used to simulate polymorphism datasets.** We depict a cartoon of a coalescent tree spanning a genomic region of $L$ bases and illustrate its contributions to our summary statistics (Section SX). For each lineage at time $t$, we add $L$ to $A(t)$ if the lineage has both European and African descendants (yellow), to $C(t)$ if all its descendants are European (blue), or to $Y(t)$ if all its descendants are African (red). We repeat this process for all marginal trees in the ARG.

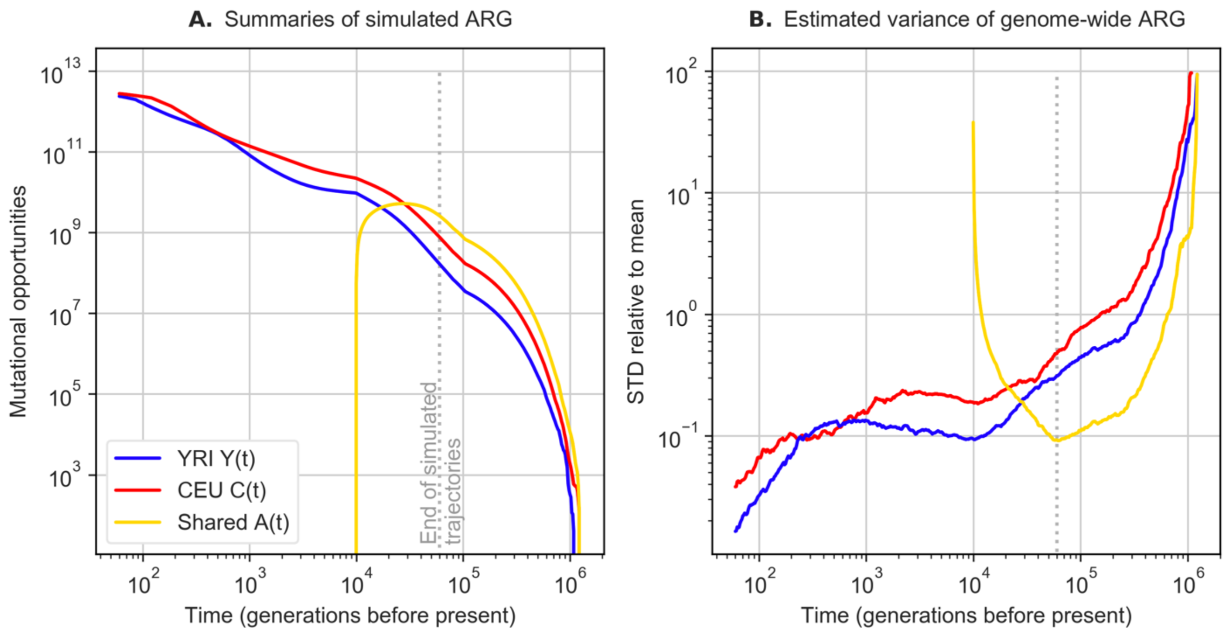


**Figure S7. Summaries of the simulated ARG as a function of time.** (A) The estimated number of mutational opportunities per generation that would result in polymorphisms that are private to the African ($Y(t)$) or European samples ($C(t)$)or shared ($A(t)$) (see Fig. S6 and SI Section 4). Almost all mutational opportunities (>99.97%) occur during the most recent 60,000 generations, which is the timeframe in which we simulate mutation rate trajectories. (B) The estimated standard deviations (relative to the mean) in our summary statistics (of mutational opportunities) if we were to simulate a sample of ARGs rather than a single one. We estimate the standard deviations by simulating an ARG for 100 genomic segments of equal size (each segment spans $3 \cdot 10^7$ bases) and estimating the variance across these segments, $\widehat{\sigma^2}$, where the standard deviation of a genomic scale simulation is then estimated as $10\sigma$. This graph suggests that variation across simulated ARGs would have a negligible effect on our results. Specifically, the standard deviation becomes substantial relative to the mean only in the distant past, in the population ancestral to contemporary Europeans and Africans, where mutation rates affecting contemporary samples were shared.

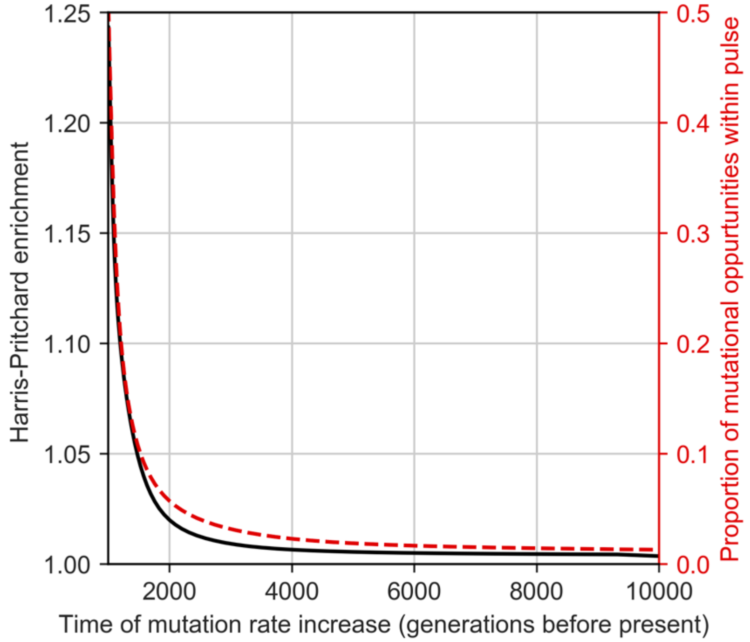


**Figure S8. The Harris-Pritchard enrichment corresponding to a pulse in mutation rate.** We show the expected enrichment (left y-axis and black curve) arising from a mutation rate pulse that starts at the time shown on the x-axis and increases the rate of one mutation type (out of the 96 types) in Europeans by 50% for a period of 1000 generations. On the right y-axis (in red), we show the proportion of the total branch length of our simulated ARG (summed across all marginal trees and weighted by the length of the segment they span) that occurs during this mutational pulse. For the purpose of this illustration, we assumed that the rate of each type of mutation is $1.25/3 \cdot 10^{-8}$ per bp per generation, except for the 50% increase in the focal type during the pulse.

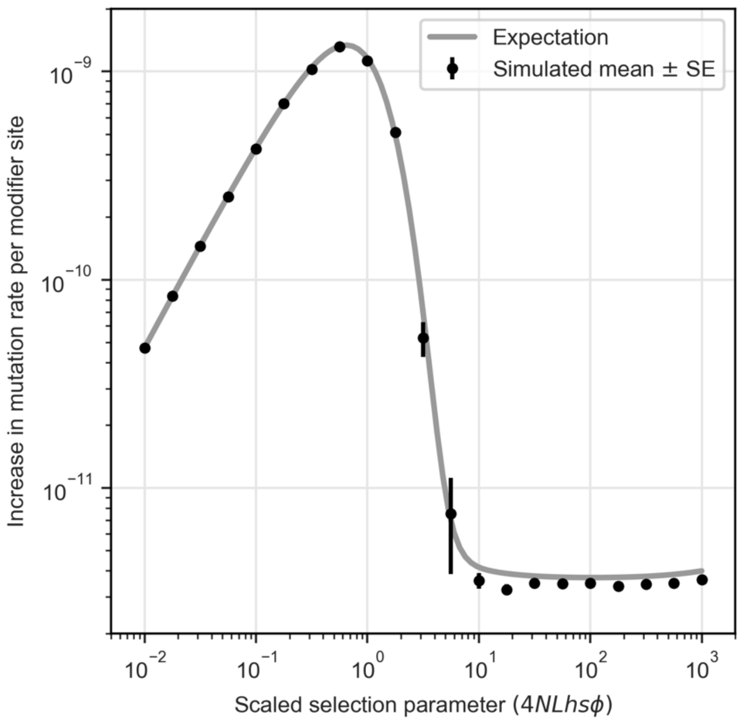


**Figure S9. Sensitivity of expected mutation rates to changes in population size.** We compare the increase in mutation rate per modifier site averaged across all simulated generations (10000 in Africans, 10000 in Europeans, and 60000 in the ancestral population) in simulations with $M = 1000$ and with changing population sizes (assuming the S&D demographic model, see Simulations) to our analytic approximation with a constant population size (Eq. 13 with $N = 1.4 \cdot 10^4$). The estimated increases and their SEs are calculated as described in *Simulations*; SEs are often too small to be seen. The average increases in simulations and in our approximation are similar throughout the range of modifier effect sizes. Specifically, we do not observe a substantial difference in the range in which $10 < 4NLhs\phi < 100$ even though such modifier sites transition from being strongly to being weakly selected during the Out-of-Africa bottleneck. The insensitivity to changes in population size changes is likely because these changes were too recent and transient (see e.g., Simons et al. 2014).

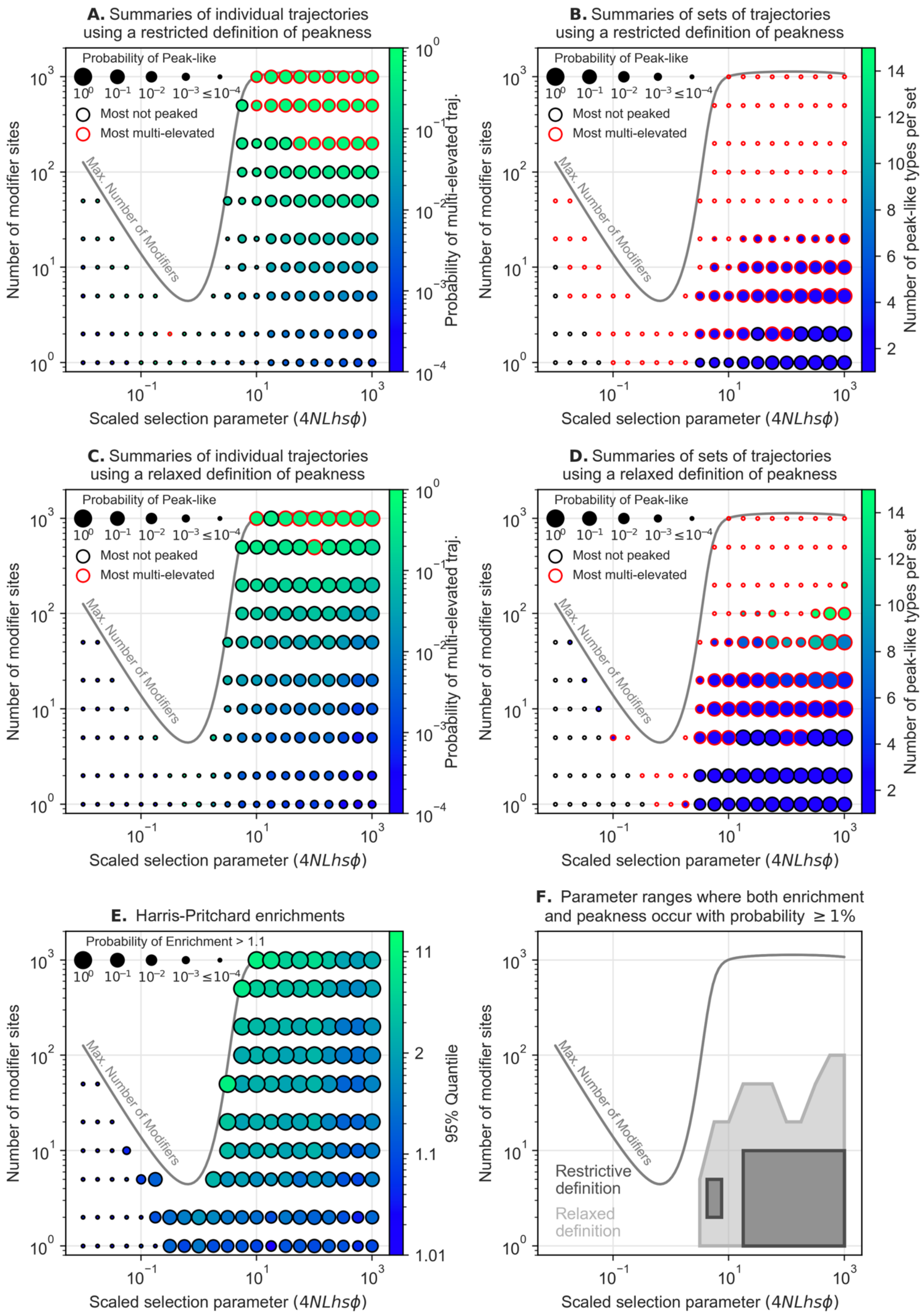

A. Summaries of individual trajectories using a restricted definition of peakness
B. Summaries of sets of trajectories using a restricted definition of peakness
C. Summaries of individual trajectories using a relaxed definition of peakness
D. Summaries of sets of trajectories using a relaxed definition of peakness
E. Harris-Pritchard enrichments
F. Parameter ranges where both enrichment and peakness occur with probability ≥ 1%
Probability of Peak-like
Probability of Enrichment > 1.1
Most not peaked
Most multi-elevated
Max. Number of Modifiers
Number of modifier sites
Scaled selection parameter (4NLhsϕ)
Probability of multi-elevated traj.
Number of peak-like types per set
95% Quantile
Restrictive definition
Relaxed definition


393

**Figure S10. Sensitivity to weak dependencies among mutator trajectories.** Our simulation approach, where we sample allele trajectories from ensembles generated from simulations with a fixed number of modifier sites ($M' = 1000$), assumes modifier sites evolve independently of each other and ignores weak dependencies that may arise between modifier sites. We examine whether our analysis is sensitive to the number of modifier sites used in simulations, $M'$, by repeating our analyses (shown in Fig. 3 and described in *Tests for variation in mutation rates between lineages*) using similarly sized ensembles generated by running 50 replicas of simulations with $M = 100$ modifier sites instead of 15 replicas with $M = 1000$ modifier sites. In (A) and (B), we repeat the analysis of individual and sets of mutation rate trajectories shown in Fig. 3A and B. In (C) and (D), we show a similar analysis as in A and B, but using a relaxed definition of *peak-like* trajectories (see Fig. S11). In panel E, we repeat the analysis of enrichment values shown in Fig. 3C. In (F), we show the parameter ranges in which both large enrichments and peak-like sets occur with probability $\geq 0.01$, similar to Fig. 4A. The results of our analysis are extremely similar to those shown in Figs. 3 and 4A, which suggests that our results are insensitive to the number of modifier sites used in simulations to generate our ensembles of trajectories, $M'$, and to weak dependencies among modifier sites.

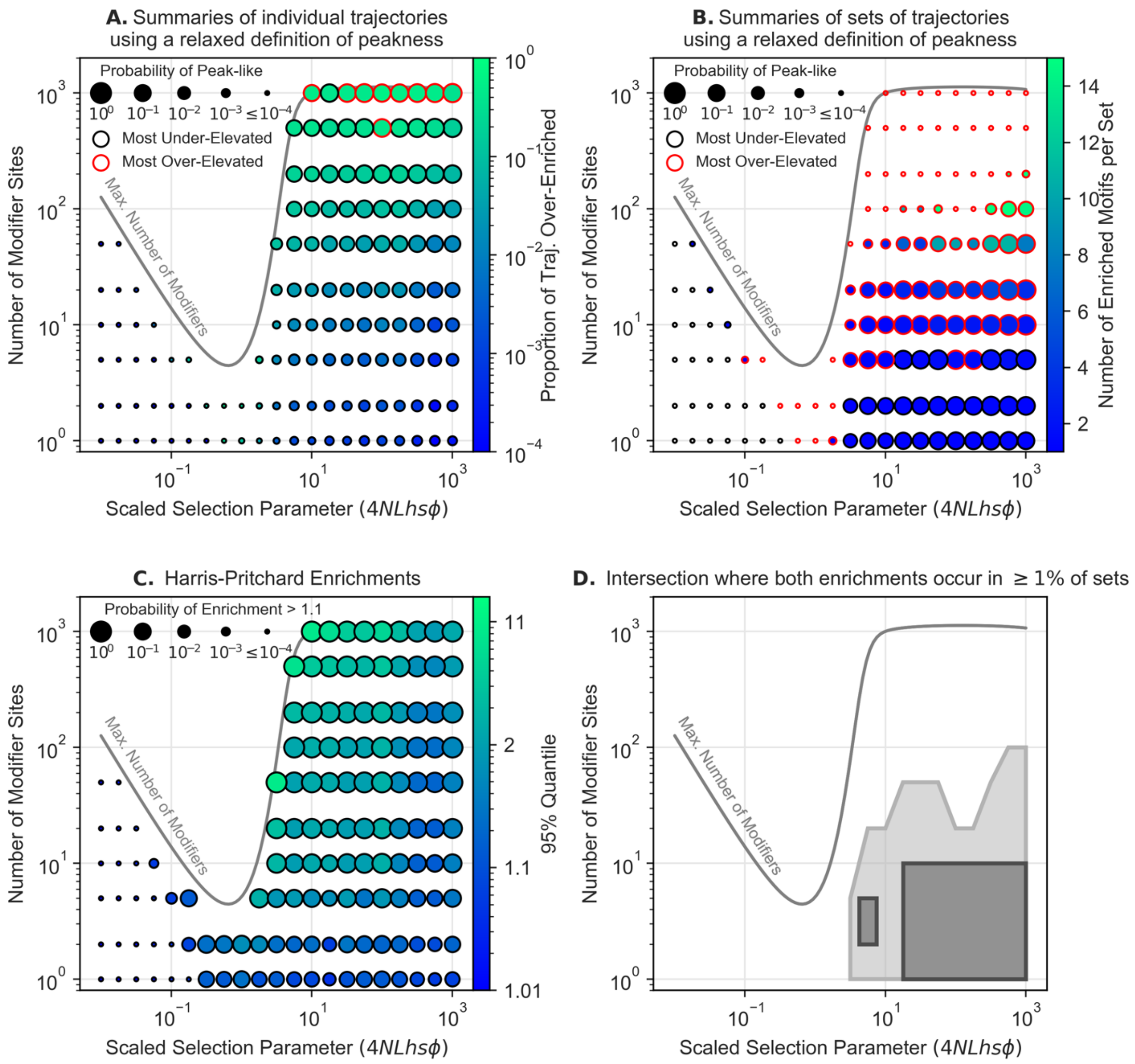


**Figure S11. Sensitivity to the definition of peak-like mutation rate trajectories**. In (A) and (B) we show the same analyses as in Fig. 3A and 3B using a more relaxed definition of peak-like trajectories (see *Tests for variation in mutation rates between lineages*). (C) and (D) are identical to Fig. 3C and 4A respectively, and are reproduced for completeness. Here, we define an *elevated* rate as exceeding expectation by more than 50% (instead of 10%), such that the *elevated* and *peaked* thresholds are the same. A trajectory is then categorized as *peak-like* if exactly one interval is *elevated*, *multi-elevated* if multiple intervals are *elevated,* or *not-peaked* if none are *elevated*. We do not change how we categorize sets of trajectories. As expected, many trajectories that were previously *multi-elevated* are now categorized as *not-peaked* or *peak-like* (A); consequently, more sets of trajectories are classified as *peak-like* (B). Enrichments are not affected by the change in definition (C). The parameter range in which both large enrichments and peak-like sets occur is expanded compared to the restrictive definition, but still restricted to $4NLhs\phi > 1$ and $M \leq 100$ (D). The discontinuities in range do not appear to be the result of sampling error alone (e.g., compare D and Fig. S10F), but we are not certain about their cause.

427

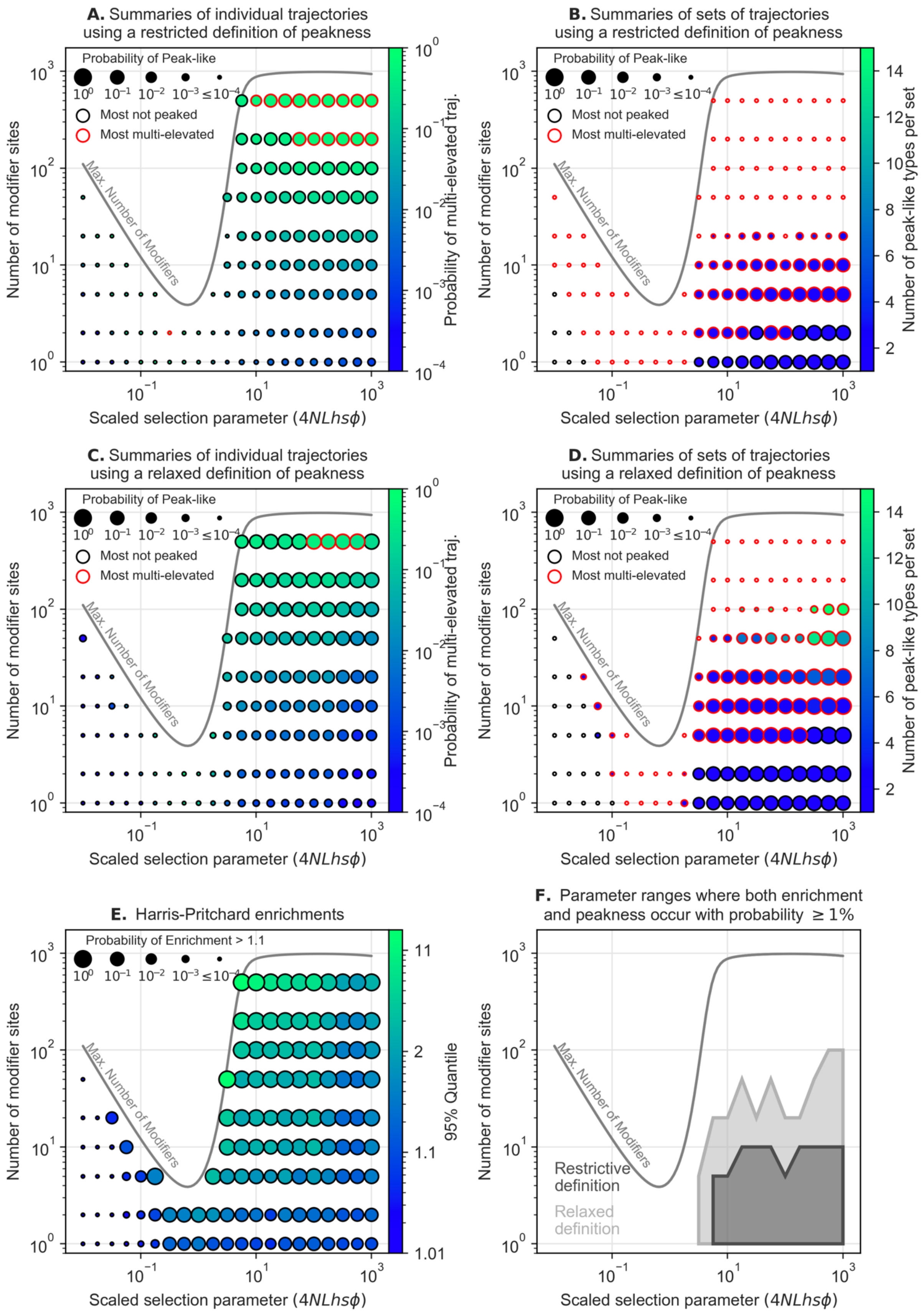

A. Summaries of individual trajectories using a restricted definition of peakness
B. Summaries of sets of trajectories using a restricted definition of peakness
C. Summaries of individual trajectories using a relaxed definition of peakness
D. Summaries of sets of trajectories using a relaxed definition of peakness
E. Harris-Pritchard enrichments
F. Parameter ranges where both enrichment and peakness occur with probability ≥ 1%
Probability of Peak-like
Most not peaked
Most multi-elevated
Probability of Enrichment > 1.1
Max. Number of Modifiers
Number of modifier sites
Scaled selection parameter (4NLhsϕ)
Probability of multi-elevated traj.
Number of peak-like types per set
95% Quantile
Restrictive definition
Relaxed definition

**Figure S12. Sensitivity to heterogeneity in rates across mutation types.** We consider a model in which the expected rate of 4 types of mutations is 5-fold higher (roughly corresponding to NCG→NTG mutations) and the rate of another 4 types is 2-fold lower than the rates corresponding to the remaining 88 types; the expected rates are chosen such that the average expected rate per base pair per generation remains $1.25 \cdot 10^{-8}$. The analyses in panels are the same as in Fig. S10 and in corresponding figures in the main text. Similar to the case with homogeneous mutation rates, *peak-like* trajectories (A and C) and sets of trajectories (B and D) are restricted to $4NLhs\phi \geq 1$, whether we use our relaxed (A and B) or restrictive (C and D) definition of peak-like trajectories. Enrichments are largely unchanged, and large enrichments are restricted to modifier sites with $4NLhs\phi \geq 1$ except for a region where $4NLhs < 1$ and $M \sim M^*(\phi)$ (E). The parameter ranges where both peak-like sets and large enrichments occur with probability $\geq 0.01$ is restricted to $4NLhs\phi \geq 1$ and $M \leq 100$ with the relaxed definition of peak-like sets, or $M \leq 10$ with the restrictive definition (F). The discontinuities in F arise from discontinuities in the parameter ranges corresponding to peak-like sets, but we are not certain about their cause.

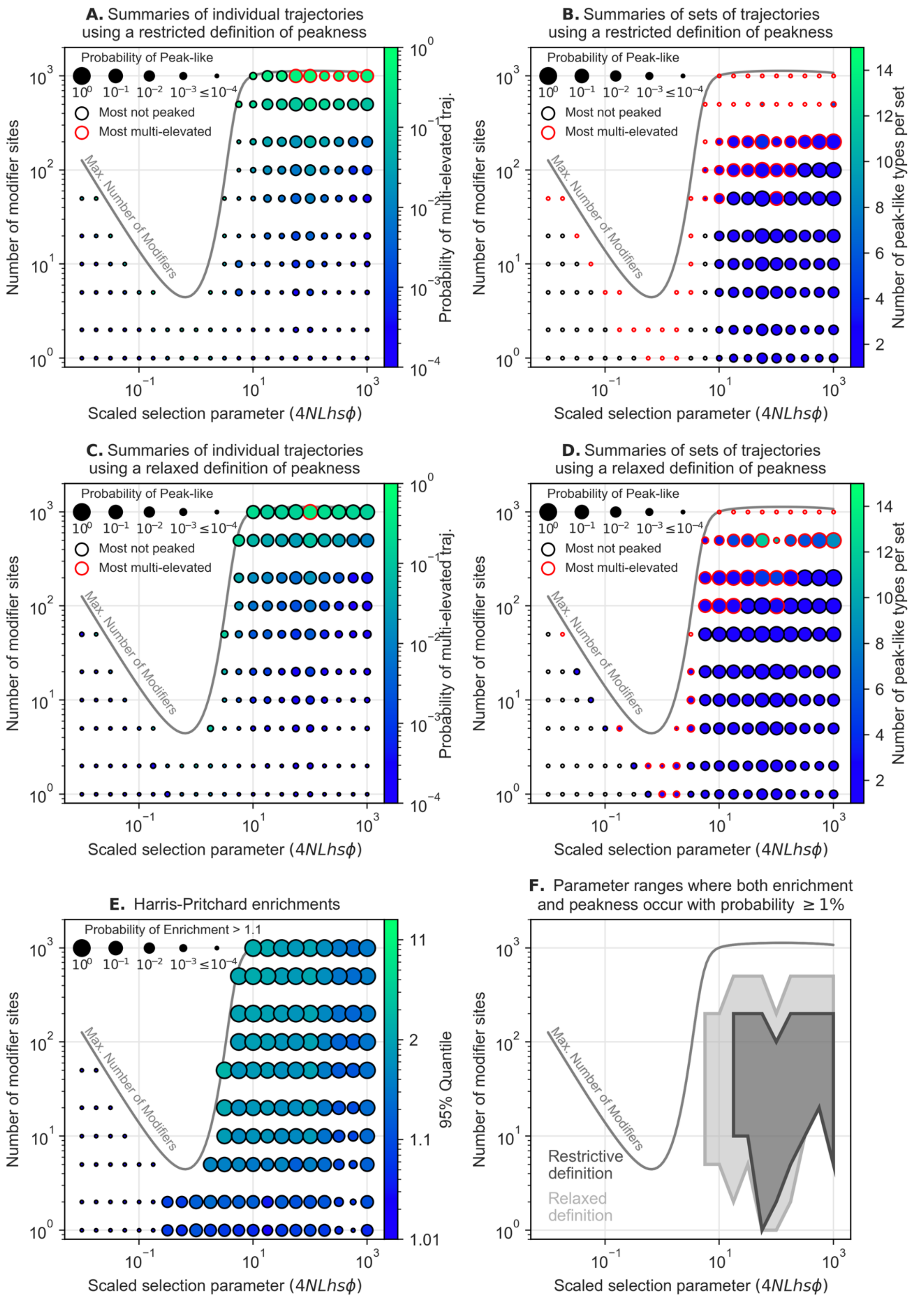


**Figure S13. Modifiers that affect the rates of several types of mutation.** In order to model correlated variation in rates of different types of mutations, we divide mutation types into

16 nonoverlapping sets of 6, with $6M$ modifier sites associated with each set. The total effect of a modifier site, $\phi$, is randomly divided among the 6 types it affects using a symmetric Dirichlet distribution (the equivalent of the uniform distribution over the (6-1)-simplex. Consequently, the scaled selection parameters associated with each modifier site remains $4NLhs\phi$ and the number of modifier sites per mutation type remains $M$. The analyses in panels are the same as in Fig. S10 and in corresponding figures in the main text. Similar to the case with modifier sites that affect one mutation type, *peak-like* trajectories and sets of trajectories are restricted to $4NLhs\phi \geq 1$ whether we use our relaxed (A and B) or restrictive (C and D) definition of peak-like trajectories. Large enrichments are restricted to modifier sites with $4NLhs\phi \geq 1$ (E). The parameter ranges where both peak-like sets and large enrichments occur with probability $\geq 0.01$ is restricted to $4NLhs\phi \geq 1$ and $M \leq 500$ with the relaxed definition of peak-like sets, or $4NLhs\phi > 10$ and $M \leq 500$ with the restrictive definition (F). The discontinuities in F arise from discontinuities in the parameter ranges corresponding to both peak-like sets and large enrichments but we are not certain about their cause.

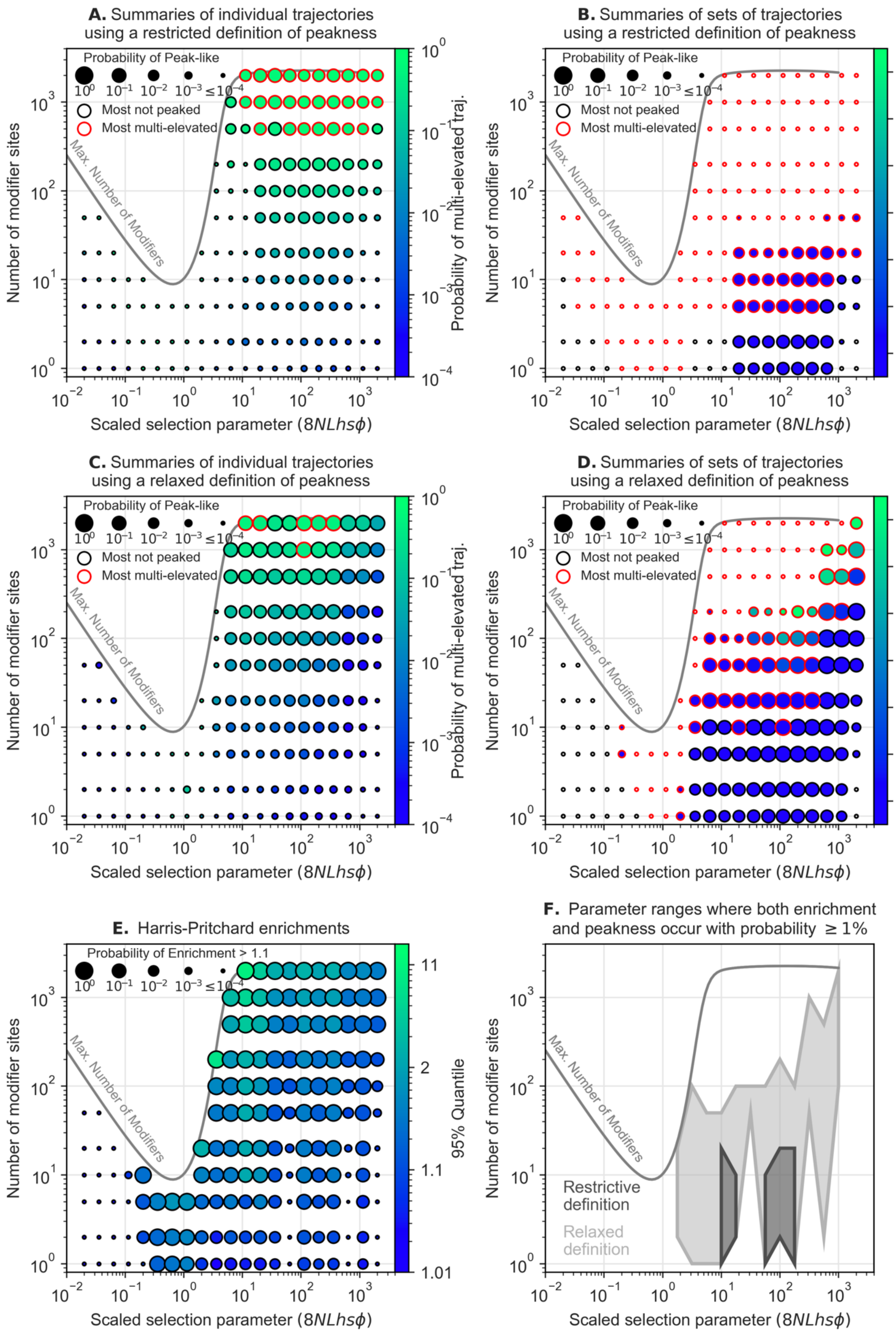

A. Summaries of individual trajectories using a restricted definition of peakness
B. Summaries of sets of trajectories using a restricted definition of peakness
C. Summaries of individual trajectories using a relaxed definition of peakness
D. Summaries of sets of trajectories using a relaxed definition of peakness
E. Harris-Pritchard enrichments
F. Parameter ranges where both enrichment and peakness occur with probability ≥ 1%
Probability of Peak-like
Probability of Enrichment > 1.1
Most not peaked
Most multi-elevated
Max. Number of Modifiers
Number of modifier sites
Scaled selection parameter (8NLhsϕ)
Probability of multi-elevated traj.
Number of peak-like types per set
95% Quantile
Restrictive definition
Relaxed definition


461

**Figure S14. A model with somatic selection against mutator alleles.** We consider a model in which selection on modifier sites arises from their effect on somatic as well as germline mutation rates, and the strength of selection due to both effects is similar (see Discussion for the consideration of other cases). Specifically, we assume that the selection coefficient of a mutator allele equals the sum of the germline and somatic coefficients, and that the somatic scaled selection coefficient equals the scaled selection parameter corresponding to the mutators effect size in our standard model. We set the scaled selection parameter (on the x-axes) to $2 \cdot 4NLhs\phi$ and recalculate $M^*(\phi)$ to reflect both sources of selection and make our results comparable to those of the standard model. Otherwise, the analyses in panels are the same as in Fig. S10 and in corresponding figures in the main text. Similar to the case without somatic effects, *peak-like* trajectories (A and C) and sets of trajectories (B and D) are restricted to $4NLhs\phi \geq 1$ whether we use our relaxed (A and B) or restrictive (C and D) definition of peak-like trajectories. Enrichments are largely unchanged and restricted to modifier sites with $4NLhs\phi \geq 0.01$ (E). The parameter ranges where both peak-like sets and large enrichments occur with probability $\geq 0.01$ is restricted to approximately $10 \leq 4NLhs\phi \leq 100$ and $M \leq 20$ with the restrictive definition of peak-like sets, or approximately $4NLhs\phi \geq 1$ and $M \leq 200$ (when $4NLhs\phi \leq 100$) or $M \leq M^*(\phi)$ (when $4NLhs\phi > 100$) with the relaxed definition (F). The discontinuities in F arise primarily from discontinuities in the parameter ranges corresponding to large enrichments, but we are not certain about their cause.

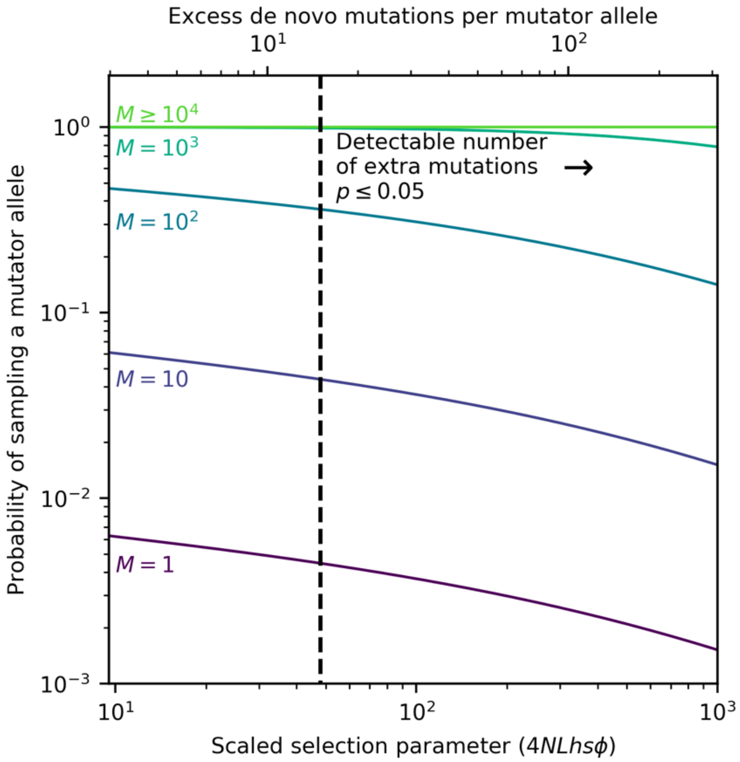


**Figure S15. Probability of sampling a mutator allele in a pedigree study with 1500 trios.** We calculate the probability using Eq. S24. When the number of modifier sites is sufficiently large ($M \gtrsim 1000$), sampling at least one mutator allele becomes very likely. The modeling assumptions and parameter values are the same as in Fig. 6.
